# Neural representations supporting generalization under continual learning

**DOI:** 10.64898/2026.01.07.698044

**Authors:** Daniel L. Kimmel, Kimberly L. Stachenfeld, Nikolaus Kriegeskorte, Stefano Fusi, C. Daniel Salzman, Daphna Shohamy

## Abstract

Abstraction and generalization are essential for flexible decision-making in novel situations. Recent work in humans and monkeys has shown how abstract variables are encoded by the representational geometry of neural population activity. However, these observations—which are typically made after learning has converged—demonstrate the *product* of abstraction, but not the *process* by which abstract knowledge is learned: how are the inputs from concrete experiences transformed into abstract knowledge, and how do neural circuits perform these operations and relay this knowledge? To address these questions, we developed a factorized model of temporal abstraction that builds on the successor representation. The model disentangles the contributions of different levels of abstract learning—from stimulus-stimulus associations to a generalizable task schema—in the form of a factorized prediction error that relates the change in relational knowledge to a predicted change in representational geometry on each trial. We fit the model to the behavior of human participants performing a context-dependent decision task during fMRI. The model captured the learning dynamics at multiple timescales, including the increasing contribution of generalization as participants transferred abstracted relational knowledge between novel task instances. In fMRI, BOLD activity in hippocampus—where, in past work, abstract knowledge was represented *after* learning—was increasingly attributed to the *acquisition* of abstract knowledge based on generalization. A similar temporal pattern was observed in entorhinal cortex, a putative source of low-dimensional structural information, and orbitofrontal cortex (OFC), which may depend on relational knowledge to represent state relationships as a cognitive map that guides choices. Indeed, individual variation in the generalization signal in OFC correlated with behavioral performance on key trials that required relational knowledge. Our findings show how the brain regions previously shown to represent abstract knowledge after learning also support the process of abstraction as it evolves from learning concrete associations to a generalizable schema. Our approach offers a computational framework for disentangling the operations driving abstract learning and probing their neural correlates in the dynamics of representational geometry.

## 1 Introduction

In a complex world, the ability to abstract a general rule, concept, or structure from a set of concrete experiences is essential for adaptive behavior. This process of abstraction allows one to generalize past experiences to novel situations by identifying their shared features. Recent work has shed light on how the brain may represent abstract knowledge in a format that supports generalization, and several theoretical accounts suggest algorithms by which this abstract knowledge may be applied in new situations (i.e., generalization) [1–5]. However, the learning process itself—how the brain extracts abstract knowledge from a set of concrete experiences—remains poorly understood.

This distinction is illuminated by recent work suggesting the brain represents abstract knowledge (i.e., the product of abstraction) in the representational geometry of neural population activity [1, 2, 6, 7]. That is, the abstracted relationships between experiences (e.g., their shared features) are represented by the distances (in neural activity space) between the neural representations of those experiences. One powerful demonstration of this hypothesis showed that when human participants learned a task’s latent structure, single-neuron hippocampal representational geometry transformed to a structured format that supported generalization [2]. Remarkably, the post-learning geometry was indistinguishable between participants who learned the latent structure via task experience and those who received verbal instructions. This finding underscores the critical distinction between the product and process of abstraction: the resulting representation of abstract knowledge may be the same for fundamentally distinct learning mechanisms.

### Learning at multiple levels of abstraction

The natural process of learning the relational structure of an environment likely unfolds continually over multiple levels of abstraction, from specific stimulus-response associations to abstract schemata that generalize to new stimuli. Here we outline the three putative levels of abstraction that we model below (see 2.2).

First, one learns concrete relationships between behavioral states (denoted below as “temporal difference (TD) learning”). For example, when going to a restaurant for the first time, one learns “after I sit down, the waiter arrives to take my order”. Given a sequence of such states, one could abstract away intervening states, for example, learning “after I sit down … the waiter (eventually) goes to the kitchen” .

Second, at a higher level of abstraction, one may learn statistical regularities that organize sets of states (denoted below as “self-regularization”). For instance, one may learn how the actions of multiple restaurant personnel are related in a general way, like “after the waiter takes my order to the kitchen, several line cooks disperse to different tasks.” Although the cooks’ specific actions may depend on the particular food order (e.g., preparing pasta), the relational structure between waiters and cooks is conserved across orders (e.g., preparing fish). That is, by generalizing the structural constraints learned in one context, one can infer the relationships between states in a different context.

Third, with further learning, one may abstract away the specific states (e.g., this waiter) to acquire a generalizable schema, allowing the transfer of abstract knowledge to novel situations with the same relational structure (denoted below as “schema transfer”). For example, after learning the general structure of a restaurant, one can quickly infer the structure of a new restaurant by mapping the novel inputs to the abstracted states of a familiar schema.

The evolution from learning at more concrete to more abstract levels potentially occurs gradually, with a time course that varies across situations and individuals. Moreover, we assume learning proceeds in parallel across different levels—learning at lower levels likely continues even after higher-level abstraction begins.

This process is unquestionably complex. Most studies attempt to mitigate this complexity by dividing the learning process into discrete learning and testing stages: learning concrete associations (and potentially the abstract structure) in one environment, then testing the generalization of abstract knowledge (if any) in a novel environment, typically without feedback to isolate the role of generalization [6, 8]. While this design affords interpretative clarity by segregating levels of abstraction to distinct phases, it obscures how learning evolves across levels. That is, what are the dynamics with which each level contributes to learning? Do multiple levels operate concurrently, and, if so, how are they distributed across the brain? Finally, how does this process vary across individuals?

In addition, most real-world environments do not provide a strong indicator, stated or implied, as to the existence of latent structure, and, in many cases, can be navigated (albeit suboptimally) without exploiting it. For example, even without a general schema for restaurants, one could enter a novel food establishment and, after some trial and error, eventually order a sandwich. In contrast, many studies of abstraction require participants to learn the task’s latent structure in order to perform reliably, including classic tasks, like Wisconsin Card Sorting and Weather Prediction [9, 10], as well as designs with a discrete generalization phase without feedback. By making abstract knowledge essential for performance, these tasks both imply (if not explicitly state) the existence of latent structure and incentivize learning at a particular level of abstraction, thereby distorting the natural evolution across levels and individual variation in the tendency to discover latent structure. These limitations would be addressed by a task in which learning the latent structure were useful (e.g., exploiting it conferred additional rewards), but not essential for adequate performance (e.g., most rewards could be obtained without learning it).

### Neural basis of abstraction and generalization

Although the neural basis of abstract learning remains poorly understood, more is known about the neural representations of abstract knowledge. A remarkable convergence of theory and experimental evidence supports a central role for the hippocampus and a network of associated regions in representing relational structure and abstracting shared features across different experiences. Hippocampal activity—measured across multiple modalities and species—encodes the relationships between behavioral states, both observable (e.g., spatial location) and hidden (e.g., latent context) [11, 12]. Moreover, recent experiments in human, monkey, and mouse have shown how latent context is encoded by the representational geometry of single-neuron population activity in a format that supports generalization [1, 2, 7]. And at least in some settings, hippocampus is necessary for adaptive behavior that depends on inferring latent states [7].

Hippocampal representations of the current environment are complemented by more general representations in entorhinal cortex (EC), where grid cell activity encodes state relationships in an abstracted, low-dimensional format that, in principle, could serve as a generalizable schema [13–16]. It has been proposed that entorhinal grid cells interact with hippocampus to map the current environment onto previously learned relational structure (i.e., generalization) [4, 5, 17].

Like hippocampus, orbitofrontal cortex (OFC) is hypothesized to represent the relationships between behaviorally relevant states [18]—often referred to as a cognitive map [19, 20]—which may be necessary for inference-guided choice [21], and hippocampal input may guide the formation of these OFC representations [22–24].

In addition, the representation of rules, categories and other abstract variables has been documented extensively in dorsolateral prefrontal cortex (dlPFC) and anterior cingulate cortex (ACC) [1, 25–29], as well as amygdala [30].

However, the above representations are typically observed after learning has converged, leaving open the question of how these representations form. In particular, what operations does the brain perform on concrete inputs (e.g., stimuli, responses, and outcomes) to learn abstract structure? And how are these operations implemented in the brain to form representations of abstract knowledge? Addressing these questions requires a behavioral task that permits expression of the natural complexity of the learning process and a framework for relating changes in abstracted knowledge to the underlying neural representations on the timescale of learning.

Here, we apply a novel computational model of abstract learning to human behavior on a novel behavioral task, and test the model’s mechanistic predictions in fMRI. Healthy participants perform a context-dependent decision-making task that permits multiple levels of abstract learning that are useful, but not essential for acquiring the vast majority of rewards. We introduce a model of temporal abstraction based on the successor representation that disentangles the contributions of each level of abstraction in the form of a factorized prediction error. Finally, the model’s factors correspond to level-specific instructions for updating the neural representation to reflect relational knowledge, which we tested in fMRI. When fit to behavior, the model reveals that humans learn at multiple levels continually, with a gradual evolution to higher-level abstraction, as participants transfer abstracted schemata between novel task instances. In fMRI, BOLD activity in hippocampus, EC, OFC, and dlPFC was increasingly dominated by teaching signals based on schema transfer. Nonetheless, in separate regions, lower levels of abstraction remained dominant throughout learning, including in hippocampus, OFC, dlPFC, and amygdala, consistent with continual learning at multiple levels. Finally, between-subject variation in the higher-level neural teaching signals predicts individual differences in behavior on key trials that require relational knowledge. Our findings support a theoretical model in which hippocampus integrates concrete sensorimotor information about external states with abstracted knowledge from current and past experiences to learn the relational structure of the environment, which downstream cortical circuits may use to represent behavioral states and direct choices. More broadly, our approach offers a computational framework for probing the formation of neural representations under continual learning.

## 2 Results

We extended a reversal-learning task from prior studies in monkey and human to permit multiple levels of abstract learning over multiple time scales [1, 2] (Figure 1). During fMRI, healthy human participants (n=39; 31 female) learned the optimal action and outcome contingencies for a set of stimuli (within-block learning). Unbeknownst to participants, the contingencies depended on two latent contexts that alternated, without cue, in blocks of trials (Figure 1b). Participants learned and exploited the context-dependent structure: a change for one stimulus was sufficient to infer the optimal actions for the other stimuli (cross-block learning). Unlike prior designs, novel stimuli were used across four consecutive sessions, thereby creating situations that required participants to generalize abstract knowledge about the task structure to these new task instances in order to perform correctly (cross-session learning).

**Fig. 1:**
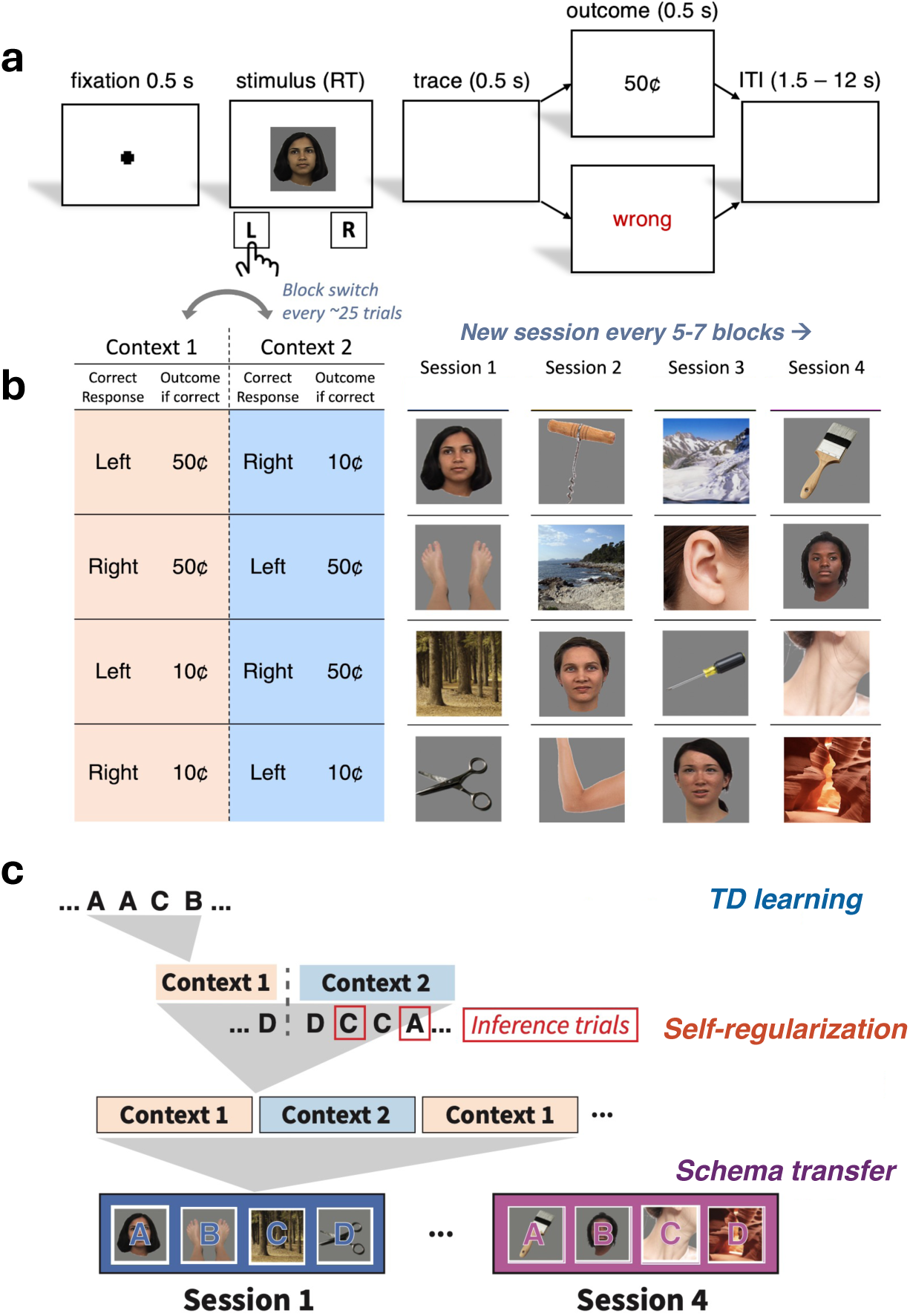
Task design. **a)** Single trial sequence. Participants were presented with single stimulus, chose the left or right action, and received deterministic feedback. (ITI = Inter-trial interval) **b)** Stimulus-action-outcome contingencies depended on latent context that alternated in 5-7 blocks of trials. Per session, 2 images from the same visual category were assigned to each “stimulus slot” (row). Novel images were used and categories were scrambled for each of 4 consecutive sessions. **c)** Task permitted continual learning across multiple time scales and levels of abstraction: local (e.g., within-block) temporal relationships between states (“TD learning”); rapid updating of optimal action on first encounter with stimuli after context switch (“inference trials”, red outlines); conserved patterns of state relationship across long time spans (i.e., between contexts; “self-regularization”); and transfer of abstracted schemata between novel task instances (i.e., sessions; “schema transfer”). Face images courtesy of Michael J. Tarr (see 4.2)

Overall, we found participants both learned in situations that did not require relational knowledge—i.e., gradually updating the correct stimulus-action associations with repetitions of the same contingencies—and also exploited the latent structure to rapidly infer the correct actions after contingencies changed between blocks and after stimuli changed between sessions. Using a computational model of the learning process, we quantified how distinct levels of abstraction contributed to learning in these different situations, and how these contributions varied over time and were represented in the brain, including explaining individual variability in behavior.

### 2.1 Behavioral results

#### 2.1.1 Inference performance

Each context switch provided participants with an opportunity to exploit the task structure. A single trial of negative feedback after the un-cued change in contingencies was sufficient to infer that the context had switched and to update the action-outcome associations for the remaining stimuli. We probed for evidence of this relational knowledge by examining performance on the first encounters with the three remaining stimuli following a context switch (excluding the first stimulus that resulted in negative feedback), which we referred to as “inference trials”. Accuracy on the three inference trials was approximately uniform (Figure 2a). We therefore defined “inference performance” as the average over the inference trials of a given block, and used this as a behavioral indicator of relational knowledge. Across all blocks and sessions, inference performance was above chance (*t*(38) = 7.75, *p <* 1.3 × 10^−9^, 1-tailed), indicating participants learned the task’s latent structure.

**Fig. 2:**
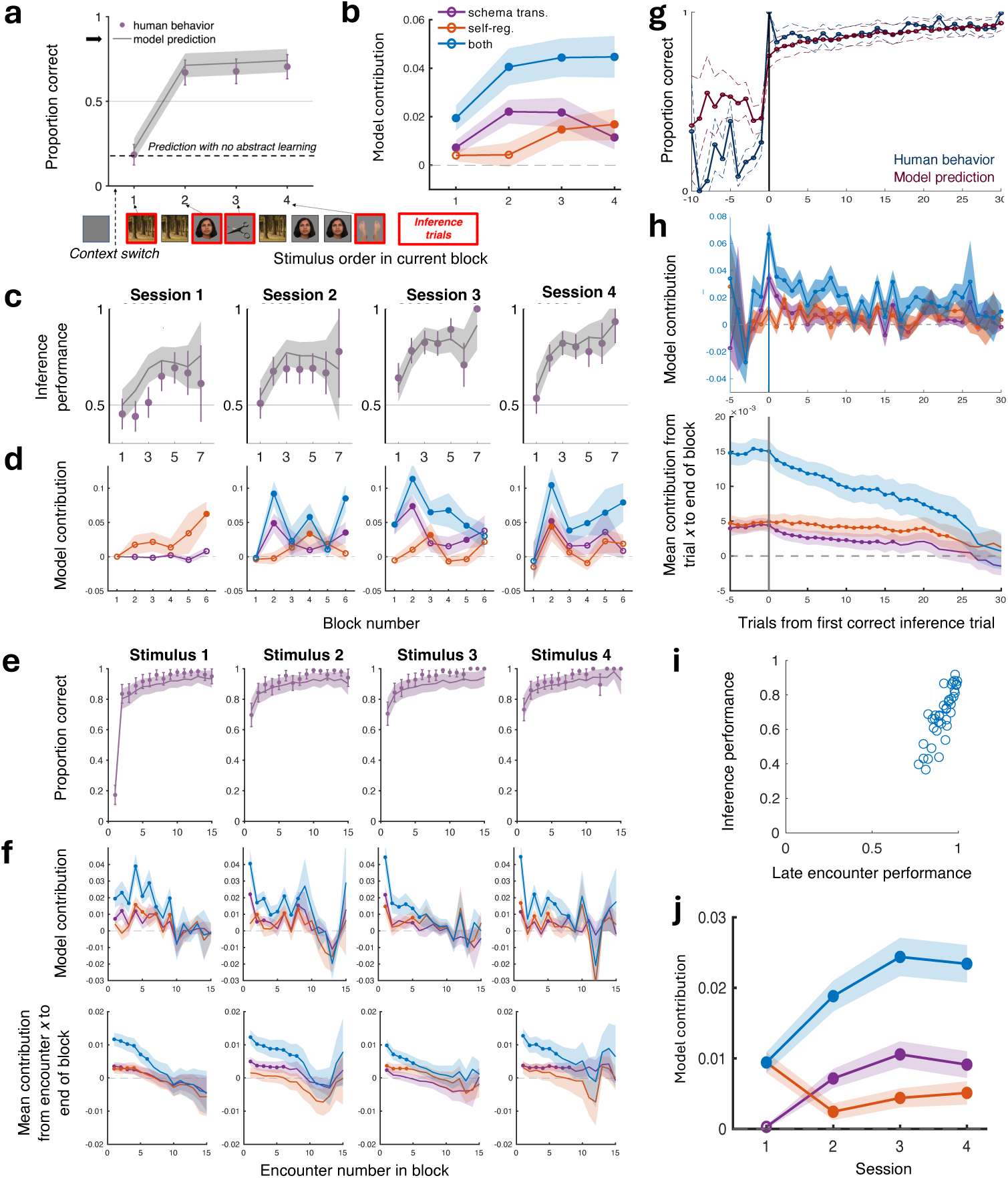
Behavioral and modeling results. **a)** Proportion correct on first encounter with stimulus after context switch as function of order stimulus was encountered in current block for human participants (circles) and gSR model prediction (line). Thick arrow shows performance at end of last block. Horizontal dashed line is prediction with no abstract learning (i.e., trial-and-error only). Chance = 0.5. Error bars and shading: ±1 std. err. prop. Example trial sequence shows assignment of stimulus order: first trial in new block is ‘Scene’, so it is assigned to ‘Stimulus 1’. ‘Face’ is first encountered on trial 3, which is assigned to ‘Stimulus 2’, etc. **b)** Contribution of model component—schema transfer (purple), self-regularization (red), or both combined (blue)—taken as the difference in log likelihood between the full model and an ablated model without the component(s), as a function of stimulus order in block. Filled circles indicate difference is significantly greater than zero (*t* -test, 1-tailed, p ¡ 0.05). Shading ±1 s.e.m. over participants. **c)** Inference performance (i.e., average accuracy over the 3 inference trials per block) as function of block, separately for each session. Markers, lines, shading, etc. as in (a). **d)** Contribution of model components on inference trials as function of block and session. Colors, shading, filled circles, etc. as in (b). **e)** Within-block learning: proportion correct vs. number of encounters with stimulus in current block, plotted separately for order stimulus was first encountered in block. Note that encounter = 1 are the same data as in (a). More than 15 encounters per block was rare and not shown. Markers, lines, shading, etc. as in (a). **f)** *Top row:* Contribution of model components vs. number of encounters with stimulus in current block. *Bottom row:* Average contribution over all encounters from the current encounter (as given by position on x-axis) to the last encounter in the block. Colors, shading, filled circles, etc. as in (b). **g)** Proportion correct as function of trials from first correct inference trial in block (*x* = 0) for participants (blue) and model predictions (red). Only trials from current block are included. More than 30 trials after first correct inference trial was rare and not shown. Dashed lines: ±1 std. err. prop. **h)** *Top row:* Contribution of model components vs. number of trials from first correct inference trial in block (*x* = 0). *Bottom row:* Average contribution over all trials from the current trial—relative to first correct inference trial in block, as given by position on x-axis—to the last trial in the block. Colors, shading, filled circles, etc. as in (b). **i)** Inference performance vs. performance on 3 or more encounters with stimulus in block for individual participants (circles). **j)** Overall contribution of model components by session. Colors, shading, filled circles, etc. as in (b). Face images courtesy of Michael J. Tarr (see 4.2)

We next examined the dynamics of inference performance over blocks and sessions. In session 1, inference remained at chance for the first several context switches. Then, beginning with block 4, inference was above chance for the remaining blocks (*t*(37) = 4.31, *p <* 5.9 × 10^−5^, 1-tailed; Figure 2c). While this behavior clearly indicated participants had acquired relational knowledge, it was less clear the extent to which this knowledge was abstracted from the specific stimuli. For example, participants could have memorized a look-up table of state sequences that was sufficient to infer the optimal actions when returning to a familiar context (e.g., “When following the Screwdriver stimulus with optimal action Left, the Forest stimulus has action Right”), where “states” were defined as the eight unique combinations of stimuli, optimal actions, and outcomes.

However, at the beginning of session 2, a look-up table strategy was not sufficient to achieve above-chance inference performance. That is, the information about specific stimuli from session 1 did not generalize to the novel stimuli of session 2. Nonetheless, we observed that participants could transfer abstract knowledge about the relationships between states defined in general terms (e.g., “Stimulus A – Left” and “Stimulus B – Right” were temporally grouped), obviating the need to learn the task structure *de novo*. We observed evidence of this higher level of abstraction in block 2 of session 2: inference performance was above chance on participants’ first-ever encounter with the states from the second context (*t*(38) = 2.95, *p <* 0.003, 1-tailed; Figure 2c). Inference performance in block 2 remained above chance in sessions 3 and 4 (*t*(38) = 5.80 and 4.40, *p <* 5.5 × 10^−7^ and 4.3 × 10^−5^, 1-tailed, respectively), consistent with generalization of an abstracted task schema to novel task instances.

While the inference performance in block 2 of session 2 depended on generalizing abstract knowledge between sessions, in principle, that knowledge could be in the form of a heuristic (e.g., given the state relationships observed in context 1, the task structure implied analogous relationships for the novel states in context 2), rather than a specific state space. In contrast, the inference trials in block 1—the very first encounters with the novel stimuli in *any* context—could only be inferred by generalizing a specific set of abstracted states to which the new stimuli were then mapped. Remarkably, block 1 inference performance was above chance in session 3 (*t*(38) = 3.09, *p <* 0.002, 1-tailed; Figure 2c) and equivocally above chance on the last inference trial in session 4 (*t*(37) = 1.66, *p <* 0.053, 1-tailed). (In general, performance in session 4 declined slightly, which we attributed to attentional fatigue.) This zero-shot learning was possible by using prior knowledge about the distribution of stimulus-action states, combined with experience with the immediately preceding stimuli, to narrow the set of available actions (e.g., “given the last two stimuli had optimal action Left, the next stimulus must be Right”).

Taken together, the inference performance in blocks 1 and 2 indicate that participants acquired *abstract* knowledge of the task structure—i.e., abstracted away from specific stimulus-action states—that generalized to novel task instances.

#### 2.1.2 Within-block learning

In contrast to inference trials, stimulus repetitions within a block permitted participants to learn the optimal action for each stimulus independently, without knowledge of the latent relationships between states. Over repeated encounters, performance improved smoothly (Figure 2e), ultimately reaching a high level of accuracy by the end of the block (89̃.0%, averaged over all blocks and participants; greater than chance: *t*(38) = 21.9, 1.8 × 10^−23^, 1-tailed ) .

We were interested in the interplay between the gradual improvement with repeated encounters and the rapid updating afforded by relational knowledge, as evidenced on inference trials. We examined performance on either side of the first correct inference trial in each block, reasoning that this point may indicate when relational knowledge was applied in a given block. While indeed performance jumped markedly after the first correct inference trial, it nonetheless continued to improve gradually for the remainder of the block (Figure 2g). This trend continued in later sessions, despite the task structure being well-learned (as evidence by inference performance), suggesting a continual role for more concrete levels of abstraction. These findings implied at least two forms of within-block learning driven by different levels of knowledge: rapid updating by applying existing relational knowledge from past blocks and incremental updating by refining task knowledge based on immediate experience in the current block. Below, we develop a behavioral model to dissect these contributions.

#### 2.1.3 Individual differences

While relational knowledge was useful, it was not essential for high overall accuracy, thereby permitting individual variation in the use of abstract learning (see 3.4). Indeed, participants differed substantially in inference performance (37 - 92%)—for which relational knowledge was essential—compared to relatively consistent performance after three or more encounters with a stimulus (77 - 99%), which did not necessarily depend on relational knowledge (Figure 2i). We exploit this dissociation below to isolate the neural correlates of behavioral variation specifically in higher levels of abstraction.

#### 2.1.4 Outcome value

In our task design, although outcome value depended on context, optimal behavior did not require learning this relationship (Figure 1b). We did not find any evidence that participants were sensitive to outcome value—such as higher accuracy or faster reaction times for the high-value stimuli—and therefore did not include outcome value in the subsequent modeling and fMRI analyses.

### 2.2 Generalized successor representation

#### 2.2.1 Model of learning at multiple levels of abstraction

The behavioral dynamics—rapid inference after context switches, zero-shot learning after session changes, incremental within-block updating—suggested learning occurred at multiple timescales and depended on multiple levels of abstraction, from concrete associations between specific stimuli to abstract schemata that generalized to novel task instances. Because learning at these levels operated throughout the experiment, they could not be isolated into discrete phases (e.g., early vs. late learning). Instead, we developed a computational model of the learning process to disentangle the independent contribution of each level on a trial-to-trial basis. Because the task’s fundamental structure was temporal (e.g., the temporal grouping of observable states defined the latent context), we built on the successor representation (SR) [31], a model of temporal abstraction previously applied to planning and navigation and shown to correlate with activity in hippocampus in both animal single neurons and human fMRI [5, 14, 31–33]. The model learned the temporal relationships between states—represented by the SR matrix—which it used to infer the current state (and thereby optimal action) given the previous state and current stimulus (Figure 3a).

**Fig. 3:**
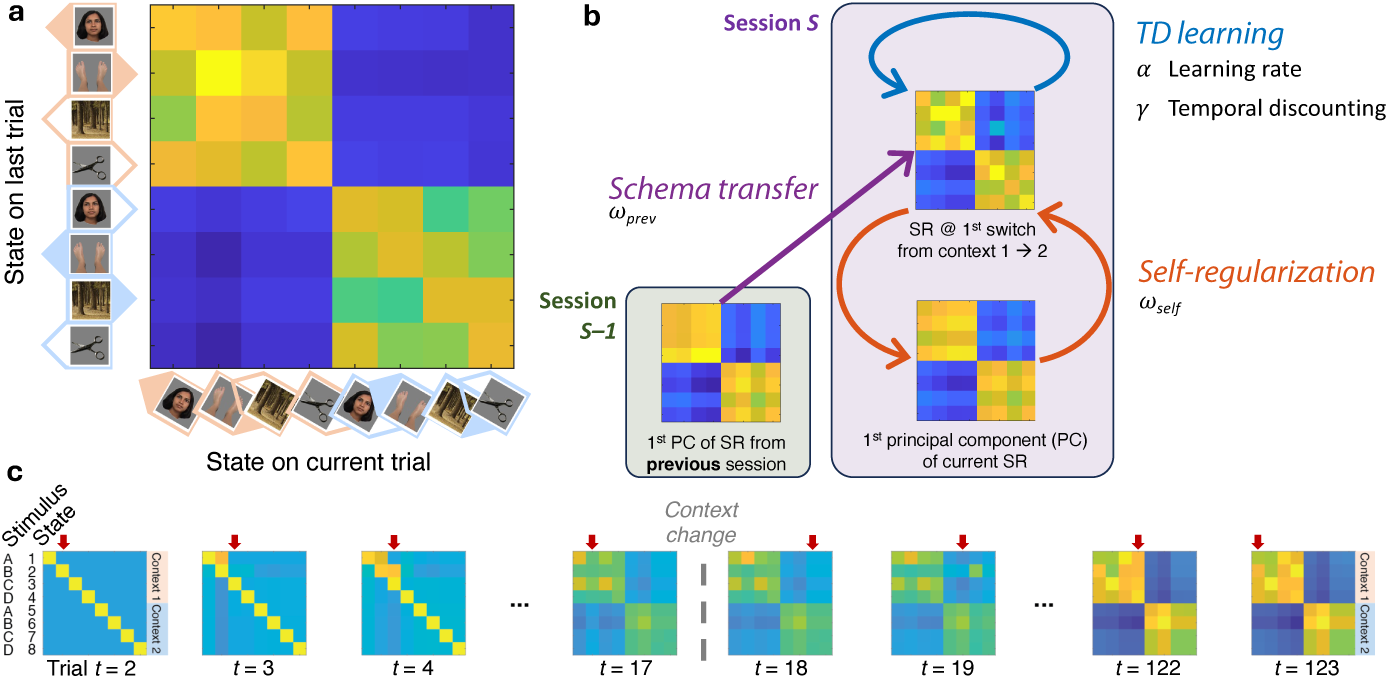
Generalized successor representation. b) gSR model update procedure … **a)** SR matrix. Elements represent long-run occupancy of state on trial *t* (*columns*) given state on trial *t −* 1 (*rows*), which agent uses to disambiguate the two possible states for the current stimulus on trial *t*. (“State” = unique combination of stimulus, optimal action, and outcome.) **b)** SR matrix is updated by factorized state prediction error with terms corresponding to level of abstraction (TD learning, self-regularization, schema transfer) and weighted by their respective free parameters *α*, *ω_self_* , and *ω_prev_* (*γ* determines temporal discounting; see main text). Low-dimensional projections of the SR matrices from the current and previous sessions are shown, as used by self-regularization and schema transfer, respectively. **c)** Trial-wise evolution of SR matrix is shown for example participant in session 3. Matrix reflects relational knowledge at time of choice on current trial. Current state is marked by downward arrow, which participant learns upon receiving feedback, irrespective of choice. On first encounter with context 2 (trial 18), SR matrix reflects relational knowledge of unobserved states, inferred via model’s higher levels of abstraction. Note example of lowest level (TD learning) learning idiosyncratic transitions between contexts (state 2 *→* 7, reflected in matrix at trial 19). Face images courtesy of Michael J. Tarr (see 4.2)

We extended the original SR model to include higher levels of abstraction, motivated by recent evidence that grid cells in entorhinal cortex encode a low-dimensional representation of the SR matrix and may supply abstract knowledge to hippocampus [4, 5, see 3.1]. Critically, our “generalized successor representation” (gSR) model disentangled the contribution of each level of abstraction in the form of a factorized state prediction error with separate terms for each level. When the model was fit to a given participant’s behavior, the trial-to-trial magnitude of each term revealed the contribution of the respective level of abstraction to the current change in the participant’s relational knowledge (as reflected in the SR matrix).

Note that, unlike the original SR model, the gSR model used reward feedback as a form of self-supervised learning—i.e., whether the recent state inference was correct—and *not* as a reinforcement signal to update a separate “reward value” vector, which the gSR model did not include. Rather, decisions were based solely on state inferences from the SR matrix, and prediction errors updated only beliefs about states, not rewards.

Here we briefly describe the model’s three prediction error terms and their corresponding levels of abstraction 1 (Figure 3b; see 4.3 for details). At the lowest level of abstraction, a temporal difference (TD) learning rule (identical to the original SR model [31]) encoded the local temporal relationships between states, with a time horizon given by the temporal discounting parameter, *γ*. Compared to the higher levels, only “TD learning” could selectively update specific state-state associations, such as the within-block refinement with repeated encounters (Figure 2e,g). As a consequence, the representation acquired by TD learning was noisy, reflecting idiosyncratic state transitions, and was constrained neither by assumptions about the relational structure nor by past experience with other task instances.

In contrast, we introduced “self-regularization” to extract structural patterns observed over all states within a single session. Self-regularization was capable of learning regularities across relationships over longer timescales, including between unobserved states, but still within the same session (thus, “self”, because it operated within-session on the current SR matrix, in contrast to across *session* schema transfer defined below). This was modeled by steering the SR matrix towards its low-dimensional projection, which may be computed by entorhinal grid cells and has been proposed as means of spectral filtering [4, 5]. In effect, self-regularization assumed that the state relationships were low-dimensional (with a dimensionality given by rank *k*, a hyperparameter), without assuming a specific structure, and is similar to other applications of low-rank regularization [34]. In the present task, self-regularization smoothed out idiosyncratic state transitions and permitted learning to generalize across contexts (e.g., given the pattern of relationships between states 1 to 4, it inferred, by symmetry, the same pattern for states 5 – 8, even before they were observed).

Finally, “schema transfer” applied the structure learned in the previous session to the current session, thereby generalizing relational knowledge without relearning. Schema transfer assumed that, across sessions, the relational structure was conserved (e.g., there were two sets of four temporally grouped states) even though the specific stimuli were novel. Schema transfer was implemented with a form of low-rank rotation and regularization that aligned the SR matrices across sessions and steered the current SR matrix toward (the aligned form of) the SR matrix learned by the end of the previous session.

At the beginning of each session, the SR matrix was initialized to an identity matrix, which meant that the SR matrix started out full-rank and was not yet representing any task structure. These conditions were contrary to the assumptions of self-regularization and schema transfer (i.e., that the task structure was low-dimensional and similar between sessions, respectively). To control for these deviations, the model weighted the terms for self-regularization and schema transfer by the variance explained by the top *k* PCs of the current and previous SR matrices, respectively. Therefore, typically for the first few trials in a session, learning was necessarily driven by TD learning, which encoded enough, albeit noisy information about state associations to inform self-regularization and schema transfer.

The TD learning, self-regularization, and schema transfer updates were weighted by the free parameters *α*, *ω_self_* , and *ω_prev_*, respectively, which, along with parameters for TD temporal discounting (*γ*) and the decision inverse temperature (*β*), were fit independently to each session for each participant. We used nested-model comparison to confirm that the model complexity (i.e., degree of freedom) added by each term significantly improved the fits. We swept through values of the rank parameters *k*—independently for self-regularization and schema transfer—to find the pair of values that applied best across all participants and sessions; we found *k* = 1 was optimal for both terms.

### 2.3 Model predictions

The gSR model reproduced human behavior at the multiple timescales and key trials discussed above, including rapid inference after context switches, both overall (2a, shaded line) and evolving over blocks (2c; see 3.6 for discrepancy in session 1, block 3); zero-shot learning in blocks 1 and 2 of later sessions (2c); and incremental within-block updating with respect to both the block onset (2e) and the first correct inference trial (2g). Note that the model predicted inference trials with high accuracy, even though they represented only a small minority of trials (°13%) and therefore had a relatively small impact on the fit likelihood. That is, the learning procedure that explained the vast majority of trials also captured behavior on a subset of key trials.

#### 2.3.1 Levels of abstraction contribute independently to learning overall

To isolate the independent contribution of a given level(s) of abstraction on any given trial, we compared the log-likelihood of the full model to a reduced model in which the corresponding term(s) was ablated. Self-regularization and schema transfer contributed significantly in all sessions they were available, although their relative roles changed over time (Figure 2j). The effect of self-regularization was greatest in session 1, when no prior experience was available for schema transfer. By session 2, the effect of self-regularization was surpassed by that of schema transfer, whose role continued to increase in session 3, as participants increasingly relied on aligning the new task instance with a familiar schema rather than re-learning the state relationships based on current experience.

We could not meaningfully test the contribution of TD learning by ablating it entirely, as it was necessary for learning the specific state-state relationships essential for early-session learning (discussed above). However, ablating only the temporal discounting component (i.e., the sub-term scaled by *γ*) confirmed that this basic form of abstraction—learning discontiguous relationships by abstracting away intervening states—played a significant, independent role.

While at least some portion of each level’s contribution was unique (i.e., could not be fully rescued by the other levels), there was some redundancy. For example, ablating *both* self-regularization *and* schema transfer caused a greater deficit in model likelihood than the sum of ablating either term alone (blue line vs. purple plus red lines, Figure 2j), indicating that when ablating only one level, the remaining level was able to partially compensate (see 3.5). The potential for interactions between the different levels of abstraction highlighted the importance of a task design that allowed each level to contribute continually—and a model to disentangle those contributions.

#### 2.3.2 Contributions to inference behavior

As with overall learning, both self-regularization and schema transfer contributed independently to the rapid updating on inference trials (Figure 2b). Moreover, as with all trials, the contribution of self-regularization decreased over sessions.

Recall that a higher level of abstraction—independent of specific stimuli—was necessary for inference in blocks 1 and 2 of later sessions (Figure 2c). Consistent with this requirement, we indeed found that schema transfer contributed significantly to zero-shot learning in block 1 in session 3, which required generalizing explicit task knowledge between sessions, and to above-chance inference performance in block 2 in sessions 2 – 4 (Figure 2d). As noted above, inference in block 2 could be achieved, in principle, by generalizing statistical regularities learned in block 1 (which the gSR modeled as self-regularization), without schema transfer. Indeed, self-regularization contributed independently to inference performance in block 2 of session 4, although its contribution to block 2 in sessions 2 and 3 was at or near zero.

While schema transfer played a dominant role overall in sessions 2 – 4, its contribution to inference performance peaked in block 2 and did not contribute significantly in the vast majority of later blocks (Figure 2d). That is, generalizing prior experience allowed the rapid acquisition of relational knowledge in a new task instance, but with time, more concrete levels of abstraction could acquire a similar degree of relational knowledge.

#### 2.3.3 Contributions to within-block learning

Unlike inference performance, learning over repeated stimulus encounters within a block did not necessarily depend on higher levels of abstraction. Indeed, schema transfer contributed significantly to only 1 – 2 additional encounters with a stimulus (after the first encounter, or inference trial; Figure 2f, top row, filled markers). The contribution of self-regularization was irregular across encounters, though was most prominent for the first (i.e., un-cued) stimulus encountered in a given block, for which inference was not possible (Figure 2f, left-most panels). Notably, the first encounter with this stimulus marked the change in contexts and therefore was an idiosyncratic state transition that TD learning might learn and for which self-regularization might have been particularly useful in suppressing.

To estimate the temporal window a given level contributed, we averaged the contributions over all encounters from a given encounter *e* to last encounter in the block; the greatest encounter *e* for which this running average was above chance defined the window’s extent. By this measure, schema transfer contributed to the first 1 – 8 encounters, self-regularization to the first 0 – 4 encounters, and both levels combined to the first 5 – 8 encounters (i.e., ablating both levels impaired the model more than ablating either alone; Figure 2f, bottom row). Nonetheless, behavioral performance continued to improve outside of these windows (Figure 2e), consistent with a conserved role of TD learning—including in later blocks and sessions—thereby underscoring how learning was continual across all levels of abstraction.

While the above analysis isolated the contribution on repeated encounters with a *given* stimulus (for which lower levels of abstraction were, in principle, sufficient), it did not account for the rapid updating in beliefs about *all* stimuli (for which higher levels of abstraction were necessary), which could have occurred prior to first encountering a given stimulus. For this, we examined the contributions relative to the first correct inference trial in the block, our indicator of the moment of rapid updating, as discussed above (Figure 2h). After this trial, the role of schema transfer decayed rapidly, dropping below chance for individual trials after 1 trial and for the running average after 16 trials (note the change from counting “encounters” with a given stimulus to “trials” with any stimulus; Figure 2h). Nonetheless, behavioral performance continued to improve after this point, again consistent with continual learning at lower levels of abstraction, which we modeled with TD learning. The role of self-regularization, while smaller and rarely significant on individual trials, was more consistent and its running average extended beyond 20 trials, as did the contribution of both levels combined. It is likely that some form of dimensionality reduction (which both self-regularization and schema transfer had the capacity to apply) was useful throughout the block to smooth out local fluctuations in state transitions due to TD learning.

Taken together, we observed continual learning across multiple levels of abstraction and across multiple timescales—within blocks, between blocks, and between sessions—throughout the duration of the task.

### 2.4 fMRI results

The evolution of inference performance across blocks and sessions indicated that participants acquired abstract knowledge of the task’s relational structure, which, in past work, was encoded by the representational geometry of single-neuron population activity [1, 2]. The gSR model suggested a mechanism by which current and past experience were integrated to update this geometry to reflect task structure. (The SR matrix specified the geometry, where each matrix element gave the distance, in neural activity space, between representations of a given pair of states.) Specifically, the model’s factorized prediction error corresponded to a set of neural instructions for how the representational geometry should change, where each term isolated the instruction derived from its respective level of abstraction.

We tested these predictions in concurrent fMRI. However, a change in geometry for multiple task conditions (corresponding to the multi-dimensional prediction error instruction) must be estimated over multiple trials, thereby precluding a trial-to-trial measure. Instead, we assumed that the *magnitude* of each model term was proportional to the *magnitude* of the BOLD activity encoding the corresponding teaching signal. By leveraging the magnitude—not the multi-dimensional geometry—we were afforded a level-specific prediction of the teaching signal that could be tested in univariate fMRI regression on *every trial* —probing the neural correlates on the timescale of learning. Moreover, because the magnitudes between terms were at most weakly correlated, we were well-positioned to disentangle the independent contribution of each term to BOLD activity within the same set of voxels.

#### 2.4.1 Hippocampus

##### Schema transfer

We began with hippocampus due to its role in representing abstract knowledge after learning, including in single-neuron activity of monkeys and humans performing a similar task [1, 2]. Across sessions, BOLD activity in left anterior hippocampus correlated increasingly more with learning attributable to schema transfer than to self-regularization and TD learning combined (i.e., schema transfer *>* (self-regularization+ TD learning)*/*2; Figure 4a, top slices). This paralleled the behavioral finding of an increasing contribution of schema transfer across sessions (Figure 2j). In addition, in left posterior hippocampus, BOLD activity on correct trials increasingly correlated more with schema transfer than TD learning (Figure 4b, top slices).

**Fig. 4:**
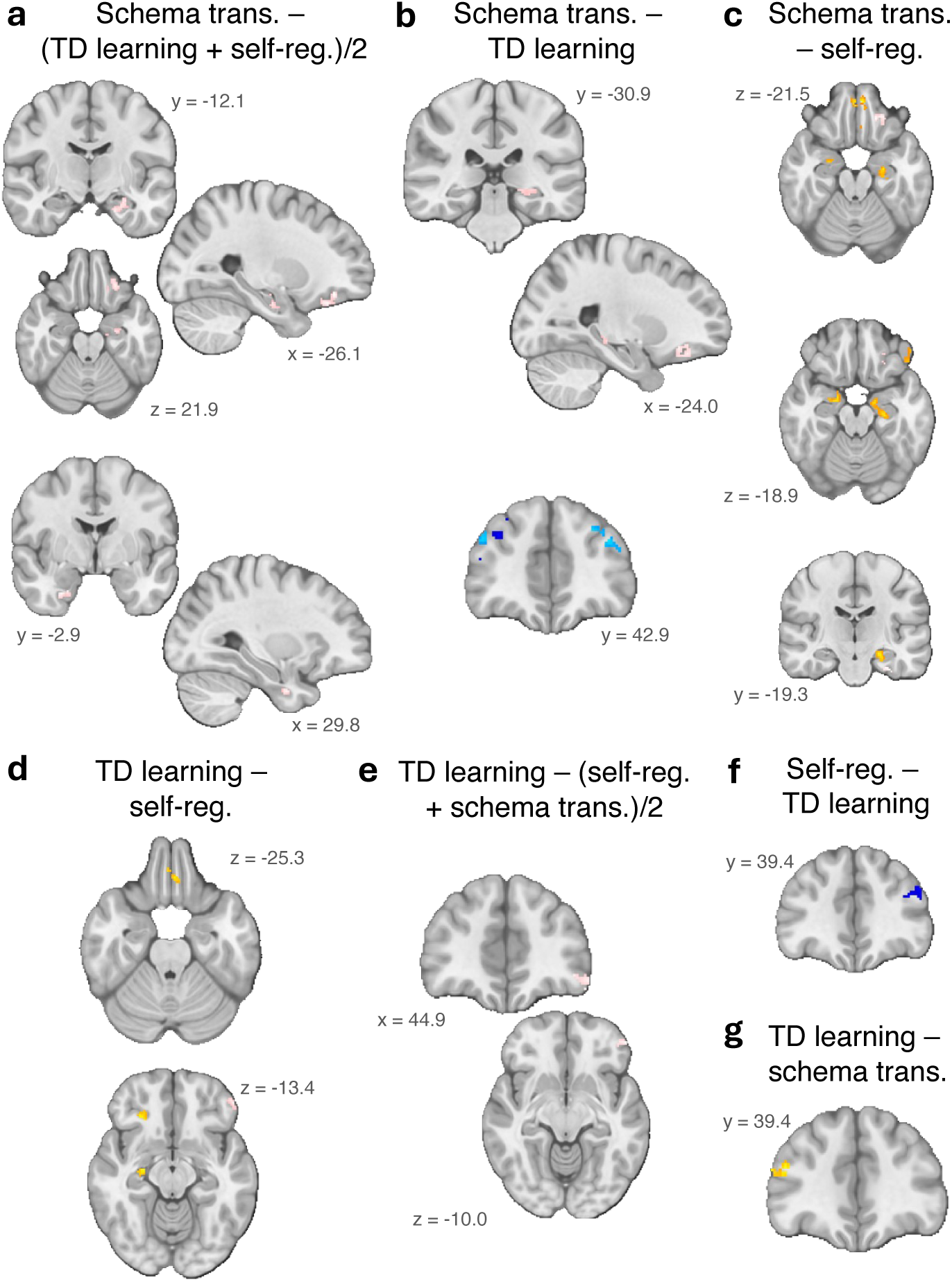
Neural teaching signals observed at all levels of abstraction across medial temporal and prefrontal cortex. **a-g)** Regions where BOLD activity was correlated more with one level of abstraction over the other level(s). Specifically, colored voxels indicate clusters of voxels (cluster extent p *<* 0.05) where the contrast A – B between two or more levels of abstraction (given by panel title) was significantly greater or less than zero (uncorrected |z-statistic*| ≥* 2.3), either on average (*red-yellow* or *dark-blue voxels*, respectively), or increasing or decreasing (*pink* or *light-blue voxels*, respectively) across sessions 2 – 4. To interpret the contrast of coefficients A – B as the BOLD activity being more modulated by level A than B, we confirmed the inequality |A*| > |*B| for each cluster. Therefore, values in (f) and (g) are sign-inverted relative to (d) and (b), respectively.

**Fig. 5:**
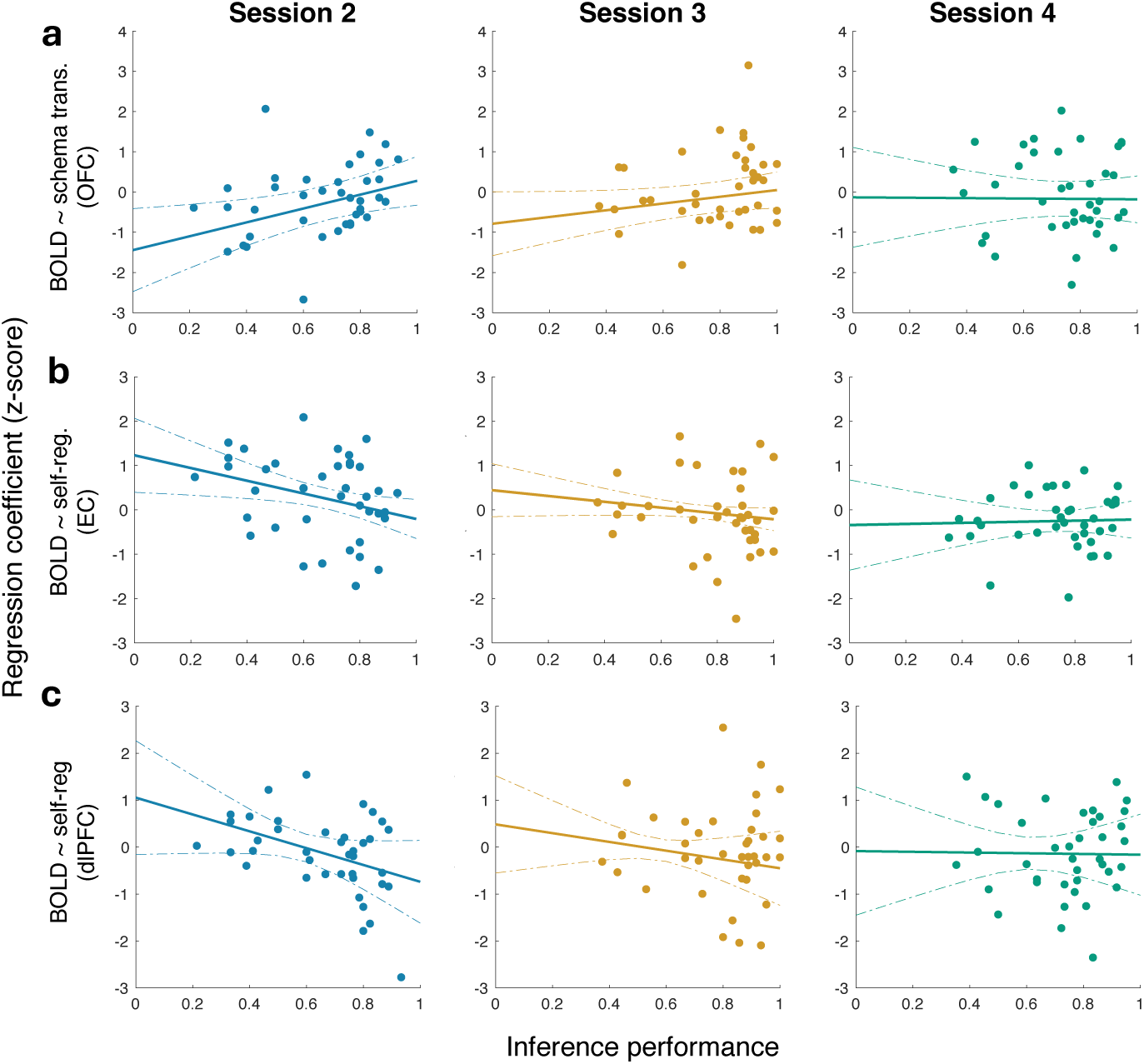
Individual variation in neural teaching signals predicts inference behavior. Participant-level regression coefficients that relate BOLD activity to a given level of abstraction (*y-axis*) are plotted against participants’ inference performance (*x-axis*) for the following specific levels of abstraction and brain regions (see main text): **a)** Schema transfer coefficient in orbitofrontal cortex (OFC) region that (at group level) prefers schema transfer over self-regularization; **b)** Self-regularization coefficient in entorhinal cortex (EC) region that increasingly prefers schema transfer over self-regularization across sessions; **c)** Self-regularization coefficient in dorsolateral prefrontal cortex (dlPFC) region that prefers (with negative sign) self-regularization over TD learning. Solid and dashed lines show the conditional predicted response and 95% CI, respectively, from linear mixed effects model (see main text).

These results are consistent with an increase in the *proportion* of the total update signal in hippocampus (i.e., component of BOLD activity correlated with the gSR prediction error summed over all levels) explained by schema transfer, and not, say, an increase in schema transfer’s contribution to behavior (i.e., increasing regressor magnitude, which would cause the regression coefficient to *decrease*) or in total brain activity (i.e., increasing overall BOLD signal, which would cause *all* coefficients to increase). (This interpretation of the increasing/decreasing trends applies throughout; for rationale, see 4.5.3.)

Because a difference between two levels of abstraction (specifically, between their corresponding regression coefficients) could be driven by either or both levels deviating from baseline, we used the phrasing “correlated more (less) with A than B” to indicate that the absolute value of coefficient A was numerically greater (less) than or equal to the absolute value of coefficient B. That is, we interpreted the difference between levels as primarily driven by level A.

In addition to examining how levels’ contributions *changed* across sessions, we also tested for regions where one level dominated the others *on average* (i.e., mean over sessions 2 – 4; session 1 was excluded because it lacked schema transfer). We found that, averaged over sessions, BOLD activity in left anterior hippocampus was correlated more with schema transfer than self-regularization (partially overlapping with the above region that changed across sessions; Figure 4c). The implications of dynamic vs. stable dominance of one level of abstraction are discussed below (see 3.3).

##### TD learning

Finally, hippocampal teaching signals were not limited to schema transfer. Averaged over sessions, BOLD activity in right anterior hippocampus was correlated more with TD learning than self-regularization (Figure 4d, bottom). That is, just as learning was explained by multiple levels of abstraction, we found that hippocampal activity was correlated with multiple teaching signals derived from those levels.

#### 2.4.2 Entorhinal cortex

##### Schema transfer

Next we considered the role of EC, which we hypothesized provided abstracted relational knowledge to hippocampus in the form of a low-dimensional projection of the current or past environment, as modeled by the self-regularization or schema transfer term, respectively. We found bilateral evidence for this hypothesis. Across sessions, BOLD activity correlated increasingly more with schema transfer than TD learning and self-regularization combined in right EC (Figure 4a, bottom slices) and than self-regularization in left EC (Figure 4c, bottom). Like in hippocampus, the temporal profile in EC mirrored the behavioral pattern: the contribution of abstracted, schema-like knowledge increased as participants transitioned from learning *de novo* about the current environment (via TD learning and self-regularization) to mapping the current task inputs onto familiar, abstracted states (via schema transfer).

#### 2.4.3 Orbitofrontal cortex

##### Schema transfer

OFC receives hippocampal input and has been proposed to represent a cognitive map of task-relevant states and features [18, 20], which it may learn in coordination with hippocampus [11, 22]. We therefore expected a similar temporal profile in OFC as hippocampus. Indeed, across sessions, BOLD activity on correct trials in the medial bank of the left medial orbital sulcus (mOS) correlated increasingly more with schema transfer than TD learning and self-regularization, both separately (Figure 4b, sagittal slice and Figure 4c, top, respectively) and combined (Figure 4a, top axial and sagittal slices).

Averaging over sessions revealed schema transfer signals in medial OFC. Specifically, BOLD activity was correlated more with schema transfer than self-regularization both in the anterior pole bilaterally (Figure 4c, top) and in a separate, more posterior region on the left midline (not shown).

##### TD learning

The contribution of TD learning changed across sessions in OFC. In left lateral OFC, BOLD activity on correct trials correlated increasingly more with TD learning than self-regularization, both separately (Figure 4d, bottom) and combined with schema transfer (Figure 4e). Unlike the behavioral parallels for the above regions, we did not find evidence of an increasing behavioral role of TD learning. Rather, it is possible that the neural systems implementing TD learning evolved over time, such as an increasing involvement of OFC.

Averaged over sessions, TD learning dominated over self-regularization in bilateral midline regions (partially overlapping with the posterior midline region that preferred schema transfer over self-regularization, discussed above; Figure 4d, top), as well as in the lateral bank of the right mOS (Figure 4d, bottom).

#### 2.4.4 Dorsolateral prefrontal cortex

##### Self-regularization

Rule learning and abstraction have long been associated with dlPFC, and dlPFC represented abstract knowledge after learning in monkeys performing a similar task [1]. We found that BOLD activity in left middle frontal gyrus (MFG) was more negatively correlated with self-regularization than TD learning on average (Figure 4f). (Unlike in other brain regions, self-regularization and schema transfer tended to be *negatively* correlated with BOLD activity in dlPFC—i.e., associated with BOLD modulations *below* baseline—and therefore BOLD activity tended to be more negatively correlated with these levels of abstraction than TD learning.) Notably, dlPFC was the only ROI we tested in which self-regularization played a dominant role. This may relate to dlPFC’s role in rule and category learning, which often depend on abstracting over the current exemplars to define an abstract state or category (like self-regularization does), as opposed to applying prior knowledge learned from dissimilar exemplars (like schema transfer).

##### Schema transfer

Compared to other regions, dlPFC showed support for all levels of abstraction. In particular, BOLD activity in bilateral posterior MFG was increasingly more negatively correlated with schema transfer than TD learning (Figure 4b, bottom). In addition, averaged over all sessions, BOLD activity in an elongated region—extending posteriorly from right SFG to MFG—was more negatively correlated with schema transfer than TD learning (MFG portion shown in Figure 4b, bottom).

##### TD learning

Finally, TD learning played a more limited but nonetheless significant role in dlPFC. Averaged over sessions, BOLD activity in posterior right MFG was more positively correlated with TD learning than schema transfer (Figure 4g).

#### 2.4.5 Amygdala

##### Schema transfer

Like hippocampus, amygdala receives extensive input from entorhinal cortex, and, in prior studies, was shown to represent abstract knowledge [30]. Averaged over sessions, we found that BOLD activity correlated more with schema transfer than self-regularization in symmetrical regions in both left and right amygdala (continuous with the anterior hippocampal regions with the same encoding, discussed above; Figure 4c, top and middle).

#### 2.4.6 Anterior cingulate cortex

As for hippocampus and dlPFC, prior single-neuron work in monkey found abstract knowledge represented in ACC [1]. However, to our surprise, we found no regions in the homologous portion of human ACC (area 24c) that were significantly correlated with the gSR prediction errors at any level of abstraction, either compared to baseline or relative to one another, and either in terms of a linear trend or averaged over sessions.

#### 2.4.7 Individual differences

We exploited the substantial individual variation in inference performance (Figure 2i) to test the link between the neural signals identified above and behavior that depended on relational knowledge. We reasoned that if the relative correlation between BOLD activity and a given level of abstraction (e.g., schema transfer) reflected a region’s specific role in the learning process (e.g., generalizing prior knowledge), then individual variation in the level’s correlation with BOLD activity would predict individual differences in inference performance. To test this link, for a given region identified above at the group level (e.g., significant contrast of abstraction level A *>* B), we related the participant-level regression coefficients for the contributing contrast (e.g., A or B) to the corresponding participants’ inference performance. In an exploratory analysis, we computed this relationship in a linear mixed-effects model with additional terms for session (fixed effect) and participant (random effect).

In OFC, we focused on the region spanning the anterior pole bilaterally in which BOLD activity was correlated more with schema transfer than self-regularization (Figure 4c, top). We found that a greater correlation between schema transfer and BOLD activity on correct trials predicted greater inference performance (inference main effect: 2.6, p = 0.03). Although this relationship appeared to lessen with increasing session number, the trend was not significant (inference-session interaction: -0.9, p = 0.12; 5a) Next we examined EC, a putative source of the low-dimensional relational knowledge necessary for the higher levels of abstraction. Focusing on the region in which BOLD activity correlated increasingly more with schema transfer than self-regularization (Figure 4c, bottom), the *less* that BOLD activity (on correct trials) was correlated with self-regularization, the *greater* was inference performance (inference main effect: -2.2, p = 0.03; inference-session interaction: 0.8, p = 0.12; 5b). That is, improved inference performance was associated with the balance of the teaching signals in EC shifting *away* from self-regularization in favor of schema transfer.

Just as relational learning depended on multiple levels of abstraction, we expected individual variation in behavior to depend on neural teaching signals beyond those of schema transfer. We therefore focused on dlPFC, where self-regularization played a dominant role. In the region in left MFG where BOLD activity was more *negatively* correlated with self-regularization than TD learning (Figure 4f), the more *negative* a participant’s correlation between self-regularization and BOLD activity, the *greater* the participant’s inference performance (inference main effect: -2.4, p = 0.03; inference-session interaction: 0.9, p = 0.11; 5c).

## 3 Discussion

While prior work showed how the brain represents abstract knowledge after learning has converged—including in the representational geometry of population activity—the cognitive operations and neural implementation of the learning process itself remained poorly understood. Our study provided a behavioral paradigm to reveal the learning process and a framework to relate changes in abstracted knowledge to the underlying neural representations on the timescale of learning. We extended a behavioral task to permit continual learning over multiple levels of abstraction—from local associations between pairs of states to abstracted schemata that generalize to novel task instances—which human participants performed over multiple timescales. We introduced a novel computational model of abstract learning that captured the human learning dynamics and, crucially, disentangled the independent contributions of each level of abstraction to the learning process on each trial. Applied to fMRI, the model’s trial-to-trial predictions—which correspond to level-specific teaching signals for updating the neural representations—revealed a network of medial temporal lobe and prefrontal systems implicated in abstract reasoning. In particular, we observed the increasing contribution of hippocampus, EC, OFC, and dlPFC to learning based on schema transfer, as participants shifted from learning associations between novel states to aligning novel states with abstracted schemata. Moreover, lower levels of abstract learning persisted in distinct regions within these and related areas, consistent with continual learning across levels of abstraction. Finally, individual variation in BOLD activity related to a region’s preferred level of abstraction predicted individual differences in inference-based behavior.

### 3.1 Hippocampus integrates across levels of abstraction

In our results, hippocampus played a central role in abstract learning. Regions in both anterior and posterior hippocampus were increasingly composed of teaching signals based on schema transfer, while separate regions in anterior hippocampus were dominated by schema transfer or TD learning throughout the task. Similarly, entorhinal BOLD activity evolved to become dominated by schema transfer, consistent with its proposed supporting role. These findings are consistent with recent theoretical work suggesting hippocampus integrates external information about current behavioral states with prior abstracted knowledge to learn the relational structure of the current environment [4, 5, 14, 19, 32].

In particular, Stachenfeld et al. [5], showed that hippocampal signals (single place cells in rodent, or fMRI voxels in human) encoded columns of the SR matrix, which described the relational structure of an already-learned environment. Furthermore, the authors showed that entorhinal grid cells encoded the eigenvectors of the SR matrix (akin to the PCA-based decomposition in the gSR model) and proposed that EC may enable a form of low-dimensional regularization to reduce noise and infer unobserved transitions during learning (though they did not implement this function in the SR model nor test its application in empirical data).

Motivated by these biological findings, we extended the traditional SR model to include higher levels of abstraction and validated the extended model in human learning data. We demonstrated how EC-based decomposition supported two, parallel learning mechanisms that explained independent components of learning. Self-regularization regularized the SR matrix toward its own low-dimensional projection, while schema transfer regularized the SR matrix toward a previously learned, abstracted schema, using the low-dimensional components to align the SR matrices from the past and present environments.

Taken together, we propose that hippocampus receives sensorimotor information about observable states (e.g., stimuli, actions, outcomes), which TD learning uses to learn an initial map of state relationships, represented by the SR matrix. Hippocampus exports this noisy and incomplete representation to EC, where grid cells decompose the SR matrix into its PCs. EC returns a low-dimensional projection of the SR matrix, toward which hippocampus steers the full-rank representation to reduce noise and infer the relationships between unobserved states (i.e., self-regularization), particularly early in learning. Over time, the entorhinal signal evolves from this self-derived target to a low-dimensional (and aligned) representation of a previous environment in which the states were superficially distinct but the abstract structure was shared (i.e., schema transfer). Finally, hippocampus integrates the outputs of these processes—TD learning, self-regularization, and schema transfer—to update its representation of the current environment.

### 3.2 Representation of abstract knowledge after learning

Hippocampus’ role in learning abstract relationships complements its previously documented role in representing abstract knowledge after learning. Our gSR model represents relational knowledge as the SR matrix, which has a natural neural implementation as a representational dissimilarity matrix (RDM), where the temporal relatedness between a pair of states (as given by the SR) corresponds to the dissimilarity between the multi-neuron or multi-voxel patterns of neural activity associated with those states [35]. The representational geometry given by the average final SR matrix in our study was consistent with the geometry observed in the hippocampal single-neuron population activity of human and monkey participants after learning a similar task: representations of task states were arranged according to their latent context, such that decoding the context for one stimulus, was sufficient to infer the context for the remaining stimuli [1, 2]. A similar geometry—one that supported generalization—was observed in hippocampus of rodents performing a simple reversal-learning task [7]. Moreover, hippocampus was necessary for a downstream state-prediction error—similar to our gSR prediction error—as well as inference-guided choices based on this error signal, consistent not only with the central role of hippocampus in representing abstract knowledge, but also with the state prediction error-based implementation of relational learning and inference proposed by the gSR model.

As in the single-neuron studies, we predict a similar representational geometry in fMRI after learning. Because our study was designed to probe the learning process, we did not collect sufficient trials at steady state to observe the final geometry, which may be particularly relevant for understanding individual differences. While we found that individual variation in the neural teaching signals was related to individual differences in inference performance, our analysis was exploratory and did not probe the geometry itself. A future dataset powered to observe the period after learning would test the hypothesis that individual variation in the model-estimated SR matrix predicts the steady-state neural geometry, which in turn predicts individual differences in inference behavior. The neural basis of these individual differences may be key to understanding the pathological variation in abstraction and generalization observed in neuropsychiatric disease.

### 3.3 Interpreting neural prediction errors

Our gSR model and theoretical framework proposes that learning at a given level of abstraction takes the form of a prediction error—computed by one or more neural circuits—that in turn updates the neural representation of relational knowledge in a downstream region. In principle, this process would account for neural activity both in the circuit computing the error and in the region receiving the error signal (which could be one and the same for a recurrent circuit). Our fMRI results cannot distinguish between these putative source and target regions, respectively. We note that, for some regions, BOLD activity was associated on average with a particular level of abstraction over all sessions, whereas in other regions the composition evolved across sessions. These temporal profiles may offer signatures of putative source and target regions, respectively, though future studies with finer-grained recording methods are required to make this distinction definitively.

In addition, prediction errors in the gSR model take the form of a vector (or matrix), which corresponds to a high-dimensional instruction for how to update the representation of one (or multiple) states—relative to the other states—in the activity space of many neurons or voxels. We circumvented the challenges of testing these multi-state, multi-voxel predictions by reducing the update to its magnitude, for which we probed in the space of single voxels. We reasoned that multi-dimensional signals projected into neural activity space would cause a mean change in population activity proportional to their magnitude. However, this proportionality does not apply to all vectors and matrices, and therefore we may be underestimating the contribution of some teaching signals to the population activity. Future work might test the specific, high-dimensional model predictions, such as the update vector magnitude *and direction* or the precise change in representational geometry. For instance, one could assume that only the representation of the present condition is updated (akin to a single row of the SR matrix) while the others remain stable, thereby obviating the need to sample representations of all task conditions for one predicted update.

### 3.4 Behavioral task that permits naturalistic learning

In contrast to prior designs, our task approximated a more naturalistic environment that permitted the learning and application of relational knowledge to occur simultaneously (i.e., not segregated into discrete phases). This allowed us to observe that humans learned continually at multiple levels of abstraction and the multiple timescales (within blocks, between blocks, and across sessions) at which these level contributed.

In addition, unlike earlier designs, learning at a particular level of abstraction was neither cued nor required. In fact, by forgoing abstract learning entirely, a participant necessarily sacrificed reward only on inference trials (i.e., 3 per block, or less than 14% of trials), while still harvesting the vast majority of rewards using trial-and-error learning alone. By placing few constraints on the learning strategy, the task allowed for the marked individual variation in abstract learning we observed (Figure 2i).

### 3.5 Extensions to the traditional SR model

#### 3.5.1 Integrating levels of abstraction

In extending the traditional SR model, the gSR not only includes multiple levels of abstraction, but also integrates these levels into a single representation of relational knowledge (i.e., the SR matrix). This integrated representation offers an advantage over alternative approaches, which typically fit multiple models—each capturing a specific level of abstraction—and either ask which model better explains the behavior (a winner-take-all solution) or attempt to mix the predictions across often dissimilar models (e.g., [8]). In contrast, the gSR model offers a more parsimonious and biologically testable mechanism in which multiple levels of abstraction operate on the same inputs and converge on a shared output space, while allowing a continuous (i.e., not winner-take-all) and smoothly time-varying contribution of each level.

#### 3.5.2 Comparing self-regularization and schema transfer

In the gSR model, the higher levels of abstraction—self-regularization and schema transfer—both regularize the full-rank SR matrix toward a low-rank projection, but the respective source of the projection differs. Self-regularization regularizes toward the *current* SR matrix, whereas schema transfer regularizes towards (an aligned transformation of) the *previous* SR matrix. Because the ideal SR matrix is conserved across sessions (given some rotation), self-regularization and schema transfer ultimately regularize towards highly similar targets by the end of learning. However, early in a given session, the targets are very different, with self-regularization operating on a sparse, high-dimensional SR matrix with only a few trial’s experience, and schema transfer steering toward a compact, low-dimensional matrix based on an entire session’s experience. Nonetheless, in the absence of one level, the other level can partially compensate. In particular, without self-regularization, schema transfer continues to provide a form of low-rank regularization because its target is itself a low-rank projection. Empirically, we find support for both aspects. Ablating either level alone impairs model performance, consistent with an independent (i.e., non-redundant) contribution of each level, while eliminating both levels impairs the model greater than the sum of either level alone, consistent with partial redundancy across levels (Figure 2b,d,f,h,j).

#### 3.5.3 Distinguishing roles of self-regularization and TD learning

Though less obvious, self-regularization also shares similarities with TD learning, both of which can learn the relationship between states that were not experienced in direct succession (e.g., learning that states 1 and 4 are related after experiencing repetition of sequences 1-2-4 and 1-3-4). However, they differ fundamentally in their mechanism and capabilities.

Self-regularization constrains associations to a limited number of patterns, in effect imposing a prior that states are related with some complexity given by the rank parameter *k* (see 4.3, eq. 7). When *k* is less than the number of states, self-regularization smooths out noisy, idiosyncratic associations, while emphasizing state relationships consistent with the expected structural pattern (defined by the evolving low-rank projection of the SR matrix). This allows self-regularization to infer relationships not only between pairs of familiar states occurring close together in time (e.g., states sharing the same context), but also over long time spans or when one or both states was never observed. For instance, after learning the state relationships in context 1, self-regularization infers the same structure for the states in context 2, even before they have been experienced.

In contrast, TD learning infers unobserved state relationships via its temporal difference term, weighted by the free parameter *γ* (see 4.3, eq. 2). With modest temporal discounting (0 *< γ <* 1), as our human participants exhibit, TD learning infers relationships between temporally proximal (though not necessarily contiguous) states. This complements self-regularization because it both operates early in learning (when the latter cannot compute a reliable low-rank projection) and permits higher resolution (i.e., higher rank) relationships (which the latter may otherwise smooth over). However, with no temporal discounting (*γ* ≈ 1), TD learning tracks *all* state transitions, not just local ones, and therefore infers unhelpful associations, such as between states from different contexts or other idiosyncratic transitions (i.e., overfitting). In contrast, self-regularization can achieve inference over long temporal spans without overfitting.

### 3.6 Model assumptions and limitations

#### 3.6.1 Choice of state space

The gSR model assumes an appropriate state space, namely that there are eight states comprised of the four stimuli paired with either of two possible optimal actions. We justify this because participants were instructed that for each stimulus, there was one of two optimal actions and that the optimal action could change over time. However, the concept that a stimulus can be in one of multiple discrete states, and switch between states abruptly, is not a trivial insight to learn, and to our knowledge, no normative theory accounts for how an agent would acquire this knowledge. Alternatively, participants could instead represent a single state per stimulus (i.e., a specific stimulus-action pair), and then update the action for that state after each context change (as in typical model-free learning). This alternative state space precludes a sustained representation of the previous stimulus-action pair and its relationships with other states, and therefore is insufficient to support the inference behavior exhibited by participants.

Nonetheless, by assuming bistable states, we are likely neglecting an important component of the human learning process—learning of the state space itself. Indeed, the assumption may account for a discrepancy between the model predictions and human choices in block 3 of session 1 (Figure 2c). In block 3, participants return to context 1 for the first time. The model retains the state definitions and relationships for context 1 that it learned in the first block, as evidenced by its above-chance inference performance. However, participants perform at chance levels on the same trials, as though they did not retain the earlier state information. It may be that when contingencies changed in block 2, participants had been representing only a single state for each stimulus, so that updating its associated action thereby discarded the original stimulus-action association (from block 1) and lead to the impaired performance observed in block 3. However, the experience of revisiting context 1’s states (in block 3) may have been sufficient to trigger a re-dimensioning of the state space to include multiple states for each stimulus (as in the gSR model), and ultimately support above-chance inference by block 4 (i.e., upon the return to context 2’s states), consistent with the gSR agent’s performance. The crucial question of how agents learn the appropriate state space would be the focus of future work.

#### 3.6.2 Aligning previous and current SR matrices

A second assumption concerns the schema transfer step, which requires aligning the current SR matrix to the SR matrix learned in the previous session. The gSR model achieves this by computing, on each trial, the rotation that aligns top *k* principal components of the two matrices. In so doing, the model assumes the alignment process is optimal, continual, and supports arbitrary rotations. In reality, human participants likely update the alignment sporadically (i.e., hypothesize a given alignment, test it for several trials, then adjust), attempt erroneous alignments (i.e., generate a non-optimal hypothesis given the observations), and limit rotations to discrete permutations (i.e., reordering the columns and rows). Future work might model how humans *learn* to align the current and previous SR matrices.

Finally, we have applied the gSR model to a limited setting in which the number of states is constant and the state relationships are of low and constant rank; defined exclusively by a single, temporal feature; and grouped into two, disjoint contexts (i.e., states were not shared by multiple contexts). Future work may extend the model to environments with higher and/or dynamic rank, multiple possible priors from which to generalize (i.e., competing schema), variable number of states across task instances, and state relationships defined by multiple, non-temporal features with several, possibly overlapping contexts.

### 3.7 Alternative models

#### 3.7.1 Hierarchical models

The gSR model learns latent structure implicitly. That is, while the SR matrix reflects the grouping of states by context, the latent variable “context” is never defined. This approach obviates the need to learn or assume the number of latent contexts or their hierarchical organization (e.g., whether groups of states are themselves grouped into higher-level contexts, and so on). In addition, the state relationships reflected in the SR matrix has a natural biological implementation consistent with observed physiology (see 3.2), whereas explicit neural representation of abstracted variables, like context, require additional assumptions or conjecture.

In contrast, a wide family of hierarchical models represent latent variables explicitly (e.g., hidden Markov models (HMMs), latent cause models, etc.) and offer complementary advantages to “flat” models like the SR. For instance, significant less memory is required to encode the one-to-many relationships between a latent state and its multiple observable states (as in hierarchical models) than to encode the many-to-many relationships between observable states (as in flat models). (Although the use of low-rank regularization in the gSR model may help mitigate these memory demands to some extent.) In addition, a hierarchical model can more easily disambiguate overlapping latent states (e.g., when a given observable state is shared by more than one latent context), whereas a flat model must encode increasingly large conjunctions of observable states, exacerbating the model’s already greater memory demands.

A recent variant of an HMM abstracts the latent structure away from the observable states by allowing a single observation to map to multiple latent states, or “clones”, thereby retaining information not only about a given observation, but also the context—or contexts—in which it was observed [36]. The model learns higher-order structure given only partial observability of the states, which is unlike the traditional SR, but may be similar in effect to the low-rank regularization of the gSR. Moreover, the model supports schema transfer by mapping new observable states to a familiar latent structure, although in the current implementation, this requires an external control to prevent new structure learning, unlike the gSR, which both transfers and updates structural knowledge concurrently without external manipulation. A future clone-based HMM adapted to multiple levels of learning may offer a smooth transition between flat and hierarchical models.

It is likely that the brain relies on both flat and hierarchical representations at different points in learning or in different brain regions. For example, early in learning, the simplicity of restricting learning to observable states may bias toward an SR-based representation. Over time, the efficiency and flexibility of a hierarchical model may shift the representation to encoding latent states explicitly. Understanding the functional implications and behavioral signatures of this putative “phase transition” would be the focus of future work.

Note that flat and hierarchical representations are not mutually exclusive, even at the same point in learning. For instance, the representational geometry that encoded multiple, abstract variables in previous work [1, 2] was consistent with an SR-like representation, as discussed above (see 3.2). However, a simple projection (i.e., linear readout) of the population activity onto a dimension encoding a given abstract variable (e.g., context) would generate an explicit representation of the variable in the downstream population, thereby achieving a contemporaneous hierarchical representation.

#### **3.7.2** Structure-transfer models and analogical reasoning

A critical component of relational learning, and likewise the gSR model, is the ability to generalize prior structural knowledge, or schemata, to novel environments and is related to the study of analogical reasoning [37, 38]. In a recent paper, Mark et al. [8], extend a typical HMM with an explicit structural prior in the form of a basis set, which must be inferred from a family of candidate structural priors and can then be applied flexibly to sets of states of arbitrary size. However, unlike the gSR, the structural priors are assumed (not learned), and structure transfer is the sole basis for learning. In contrast, the gSR model learns the structural prior *de novo* and flexibly integrates over multiple levels of abstraction (i.e., not restricted to structure transfer). In future work, a hybrid model might synthesize the advantages of the two model classes.

## 4 Methods

### 4.1 Participants

A total of 46 participants (11 M, 35 F) between the ages of 18–35 were recruited from the Columbia University community. Participants were right-handed, had normal or corrected-to-normal vision, took no psychiatric or sedating medication at the time of the experiment, had no diagnosis of psychological, neurological, cognitive, or emotional disorders, and had no contraindications for magnetic resonance imaging (including but not limited to pregnancy, claustrophobia, history of high exposure to metallic particles, and incompatible implants). One participant could not tolerate the imaging procedure and aborted the experiment before beginning the behavioral task. Six participants were excluded for inattention during the behavioral task (see 4.6). The remaining 39 (8 M, 31 F) participants had a mean age of 25.1 years (range 18–35) and were included in the reported sample. No statistical method was used to predetermine sample size. Written informed consent was obtained at the beginning of the experiment, and all experimental procedures were approved by the Columbia University Institutional Review Board.

### 4.2 Behavioral task

We extended a reversal-learning task from prior studies in monkey and human to include multiple levels of abstract learning [1, 2]. In summary, participants learned the optimal action and outcome contingencies for a set of stimuli (within-block learning). Unbeknownst to participants, the contingencies depended on two latent contexts that alternated in blocks of trials. The context-dependent structure could be exploited to maximize reward: a change in one stimulus was sufficient to update responses for the other stimuli (cross-block learning). Across sessions, novel stimuli were used, permitting generalization of the task structure to these new instances of the task (cross-session learning).

The entire behavioral task was performed during fMRI acquisition (see 4.4). Stimulus presentation and behavioral response monitoring was implemented in custom code written for the *PsychoPy* package and running on a Macbook Pro laptop (Apple, Cupertino, CA). Stimuli were projected onto a screen positioned at one end of the scanner bore. Participants viewed the screen while lying supine in the bore via a mirror angled at 90°and positioned just above their eyes. The projected background was set to black and text was presented in white, unless otherwise noted. Behavioral responses were collected via a pair of button boxes, one in either hand.

The beginning of a trial was cued with the onset of a white central point (0.5 s), followed by the presentation of a single stimulus (Figure 1a). The participant then rendered their choice, “Left” or “Right”, indicated by a button press with the respective left or right thumb. The stimulus presentation terminated either after the choice was rendered (i.e., reaction time, RT) or 1.6 s, whichever was sooner. If the participant did not respond in time, or pressed both buttons simultaneously, an error message was displayed for 1 s (i.e., until what would have been the end of the outcome epoch had a valid response been provided) and the trial was scored as invalid. Otherwise, a blank screen was presented (0.5 s) followed by the outcome (0.5 s), which was deterministic: when correct, the amount of reward was displayed in magenta numerals (RGB = [1, 0, 1]); when incorrect, the word “Wrong” was displayed in red (RGB = [1, 0, 0]). The outcome was followed by the inter-trial interval (blank screen) composed of a fixed interval (1.55 s) and a random interval drawn from a truncated exponential distribution (mean = 3.1 s; max = 10.85 s), which combined to a nominal mean of 4.65 s (i.e., 3 TRs) and range of 1.55 to 12.4 s (1 - 8 TRs), appended to which was any unused portion of the stimulus presentation (i.e., 1.6 s minus the RT), such that the interval between trials did not depend on the participant’s reaction time.

Stimuli belonged to one of four visual categories (faces, body parts, tools, and scenes). Except for scene images, stimuli were isolated from any background and placed on a neutral gray background. For a given session (see below), two unique stimuli were used for each category (i.e., total of 8 unique stimuli per session), and each stimulus category was mapped to a single “stimulus slot” that governed the the response and outcome contingencies for the pair of stimuli from that category (Figure 1b). (Once the stimulus slot was assigned for a given trial, the specific stimulus was assigned uniformly, p = 0.5, over the pair of available stimuli.) The stimuli within a category were functionally equivalent, and, from the earliest trials, participants responded indistinguishably to the two images from a given slot (e.g., once updating their response for one of the faces, they applied the same updated response on their first encounter with the second face). That is, participants instinctively generalized across images from the same visual category; therefore we analyzed the data with respect to the stimulus slot (i.e., in effect, coding the two images as a single stimulus). For simplicity, we used the term “stimulus” to refer to a particular stimulus slot (i.e., either of two images) and “image” to refer to a unique image.

Stimuli were obtained from publicly available image databases. Face images courtesy of Michael J. Tarr, Carnegie Mellon University, http://www.tarrlab.org/, with funding provided by NSF award 0339122.

Each unique combination of the three observable variables—i.e., stimulus, action, outcome contingency—defined one of eight conditions, or “states” (Figure 1b). The contingencies for the four stimuli remained stable for a block of trials. Block duration was drawn from a truncated exponential distribution (mean = 15; max = 45) and added to a fixed 10 trials, for a combined nominal mean of 25 trials and range of 10 to 55 trials. In alternating blocks, the contingencies switched back-and-forth, *without cue*, between two sets of contingencies. Each set corresponded to the latent variable, “context”, defined only by the temporal grouping of the contingencies. The conjunction of context and stimulus was sufficient to infer the optimal action and outcome value, even on the first encounter with a stimulus after a block switch.

Within a given block, the sequence of stimuli were drawn as follows. On the first trial of the block, the stimulus was selected with uniform probability (p = 0.25) over the set of four stimuli. On subsequent trials in a block, the stimulus depended on the stimulus in the previous trial. Specifically, within each context, the four stimuli A, B, C, D were grouped into two pairs, or two “subcontexts”. In context 1, stimuli were grouped into subcontexts composed of A, D and B, C, and in context 2, into sub-contexts of A, B and C, D. The current subcontext was biased (p = 0.7) to be the same subcontext as on the previous trial, and otherwise switched to the other subcontext. Once subcontext was determined, the stimulus was uniformly selected (p = 0.5) over the two available stimuli for that subcontext. This system for sequence generation created short, alternating sub-blocks containing pairs of stimuli, thereby defining an additional, orthogonal latent variable, subcontext, that had no instrumental value (unlike the context variable that determined the contingencies). We did not exploit this additional task feature in the present analyses.

Participants completed four consecutive task “sessions” without exiting the MRI scanner. Each session contained a stable set of stimuli in alternating blocks of contexts 1 and 2 for a total of 15 minutes of scanning time (mean = 5 blocks; range = 3 – 8 blocks). The session was divided into two, equal runs (7.5 minutes per run), between which the task was briefly paused with at least 10 trials remaining in the present block. At the end of the second run, participants were told how much reward they had earned in that session followed by a brief pause.

Across sessions, novel images were used for each visual category, and the mapping of visual category to stimulus slot was shuffled systematically, thereby eliminating the utility of specific image or category information from prior sessions, but permitting generalization of the abstract structure: four stimulus slots with specified action and outcome contingencies that depended on latent context.

After completing the tasks, the participant received the earnings from one of the four sessions, selected at random, as a bonus.

#### Participant instructions and practice

Immediately prior to the experiment, participants were instructed on the task. They were told to learn the correct response (Left or Right) to each of a series of pictures and would receive reward of 10 or 50 cents, depending on the picture, for correct responses. They were told that “every once in a while the correct response will change,” and they must learn the new response, and that the amount of money earned may also change. No additional information about the task’s latent structure was provided. They were told they would complete a total of four sessions, but not that the pictures would change between sessions nor what, if anything, about the task structure would be maintained between sessions. They were told they would receive bonus money based on rewards earned from one session selected at random.

After receiving the instructions and before entering the MRI scanner, participants performed a brief practice version of the task that included two fractal stimuli (i.e., unrelated to the visual categories in the experimental task) with opposite optimal actions that remained static throughout the practice round (i.e., contingencies did not change). As in the experimental task, a brief pause was included midway. A small number of correct trials was required to advance to the experimental task.

### 4.3 Generalized successor representation

#### 4.3.1 Overview

The successor representation (SR) is a model of temporal abstraction using a temporal difference (TD) algorithm to learn the relationships between behavioral states ([31]). We extended the SR model to include multiple levels of learning, including the capacity to generalize state relationships between distinct task instances, that we call the generalized SR (gSR) model.

We applied the gSR model to the present experimental task. On each trial *t*, the agent observes stimulus *o_t_* and must select an action *a_t_* ∈ {*R, L*}. If *a_t_*is equal to the optimal action *a*^∗^, the agent will receive a positive reward; otherwise the agent will receive 0 reward. Given four stimuli and two optimal actions, there are a total of eight possible states *s_t_* = (*o_t_*, *a_t_*^*^). Given stimulus *s_t_*|*o_t_* ∈ {(*o_t_*, *a_t_*^*^ = *R*, the agent must disambiguate between two possible states: (*o_t_*, *a_t_*^*^), (*o_t_*, *a_t_*^*^ = *L*)} based on the prior state and the temporal structure of the task.

Our implementation of the SR differs from prior uses in several key ways:

- Traditionally, the SR is used to infer the future state(s) *s*^′^ based on the current state *s_t_*. Here, we use the SR to infer the current state *s*^′^ based on the prior state *s_t_*_−1_. To emphasize this distinction, on the current trial *t*, we index the rows and columns of the SR matrix *M_t_* with *s_t_*_−1_ and *s_t_*, respectively. When the current state is uncertain (i.e., prior to the choice), we refer to the current state as *s*^′^, which still indexes the columns of *M_t_*.
- After rendering a choice, the agent receives deterministic feedback (reward for correct, null for incorrect) that is sufficient to completely resolve the current state from the uncertain case *s*^′^ to the known case *s_t_*. The agent uses this feedback to update the appropriate row and column *M_t_*(*s_t_*_−1_, *s_t_*) given the *veridical* state *s_t_*, *not* the state given by the agent’s choice. As such, reward in this case serves as a label (correct, incorrect) for self-supervised learning, not as a value signal for reinforcement learning, as in traditional uses of the SR.
- The SR matrix *M_t_* is updated via several additive terms that factorize the state prediction error (PE) into different levels of abstract learning:
  1. **Temporal difference (TD) PE**. This term, which is the only term shared by the gSR and traditional SR models, uses a TD learning rule to encode the temporal structure of state transitions but does not impose structured relational constraints. The TD PE models the lowest level of abstract learning that captures short-range relational dependencies.
  2. **Self-regularization PE** applies low-rank regularization based on the current SR matrix *M_t_*, hence is a form of self-regularization that forces relational knowledge to be represented in a simplified form that spans multiple contexts. Self-regularization extends learning over longer time spans, extracting structured relationships across multiple contexts under a dimensional constraint.
  3. **Schema transfer PE** applies low-rank rotation and regularization to align learned relational structures across tasks instances, facilitating transfer without relearning. Specifically, the term regularizes the current SR matrix *M_t_* toward (the aligned form of) the SR matrix *M^prev^* that the agent learned in the previous task session, which shared the same temporal structure but used different stimuli. The schema PE models the highest level of abstraction, in which the agent abstracts a relational schema, enabling the application of learned structure to new but structurally identical settings.

Each term is scaled by a free parameter:

*α*: TD learning

*ω_self_* : Self-regularization

*ω_prev_*: Schema transfer

#### 4.3.2 Procedure

The model is implemented in MATLAB (Mathworks, Natick, MA, USA) using custom code. On each trial, the following steps are performed:

- Given the observed stimulus *o_t_* on the current trial *t*, consider the two possible states (*o_t_, a*^∗^ = *R*) and (*o_t_, a*^∗^ = *L*), which for simplicity we will denote as *s_t_^R^* and *s_t_^L^*, respectively.
- Choose action *R* with probability *p*(*s_t_^R^*) given by the difference in state occupancy of the two possible states for the current stimulus *o_t_* as taken from the row of the SR matrix *M* corresponding to the preceding state *s_t_*_−1_, transformed by the softmax function with inverse temperature *β* ∈ [0, ∞] :

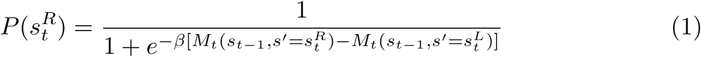 *Analogy to Q-learning:* Assuming that producing the correct action yields 1 unit of reward and the incorrect action yields 0, the value *Q* of each possible state is proportional to the probability that the agent has inferred the state correctly, that is *Q*(*s_t_^R^*) = *M_t_*(*s*_*t*−1_, *s*′ = *s_t_^R^*. As such, the probability of choosing *R* is proportional to the difference in value of the two possible states, *Q*(*s_t_^R^*) − *Q*(*s_t_^L^*. This formulation is analogous to the traditional use of the SR for reward-guided reinforcement learning, in which the Q-value for each stimulus-action pair is taken as the product of the SR and a learned reward vector, and then choice is determined by the relative Q-values of the two possible pairs for the current stimulus.
- Update row *M̂* (*s_t_*_−1_, :) for all states *s*^′^ given prior state *s_t_*_−1_ and the veridical current state *s_t_* (which is resolved for the agent following the outcome):

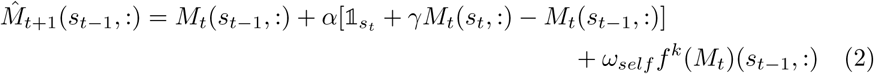

where *α* ∈ [0, 1] is the learning rate, *γ* ∈ [0, 1] is the temporal discounting factor (when *γ* = 0, *M* reduces to the one-step Markovian transition matrix), and *ω_self_*∈ [0, ∞] scales the self-regularization function *f* (see below). The one-hot row vector 1 has value 1 for the current state *s_t_* and 0 otherwise. Note that the terms inside the brackets corresponds to the TD PE (above) and is analogous to the traditional TD learning rule for value functions, except that the reward prediction error is replaced by a successor prediction error. The term scaled by *ω_self_* corresponds to the self-regularization PE (above).
- Update all rows of *M* , combining the just-updated row *M*^^^*_t_*_+1_ and the schema PE (above):

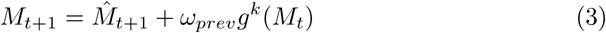

where *ω_prev_* ∈ [0, ∞] scales the schema transfer function *g* (see below).

We assume the SR matrix *M* is initialized to the identity matrix, that is *M* (*s_t_*_−1_, *s*^′^) = 1 if *s_t_*_−1_ = *s*^′^, and *M* (*s_t_*_−1_, *s*^′^) = 0 if *s_t_*_−1_ ≠ *s*^′^. This initialization implies that each state will necessarily predict only itself. (Note that one could initialize *M* with additional assumptions, e.g., that states sharing the same stimulus *o* are anti-correlated.)

In sum, the model has five free parameters: *α*, *γ*, *ω_self_* , and *ω_prev_*for learning, and *β* for choice.

#### 4.3.3 Regularization Details

Regularization by *f^k^*(*M_t_*) and *g^k^*(*M_t_*) consists of two aspects that achieve, respectively: a) rank-based self-regularization using the present SR matrix *M_t_* and b) schema transfer using *M^prev^*, which was learned by the agent on a prior instance of the task. Both aspects rely on the eigenvectors of *M* , which has been proposed to be computed by grid cells in entorhinal cortex and input to the place cells in hippocampus, which in turn represent columns of the SR matrix ([5]). On each trial *t*, prior to updating the SR matrix, we find the principal components (PCs) *U* of the SR matrix and their corresponding eigenvalues *V* using principal component analysis (PCA).

In subsequent steps, we will weight the contribution of the regularization term by the variance explained by the top *k* PCs of the relevant matrix, *M_t_* for self-regularization *f* or *M^prev^* for schema transfer *g*. Variance explained is computed from the eigenvalues *V* as:

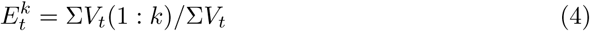

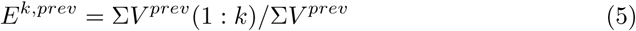

For the subsequent steps:

- The columns of *M_t_* and *M^prev^* are first mean-centered.
- Let *U_k_* and *U_k_^prev^* refer to the top *k* PCs of *U* and *U^prev^*, respectively:

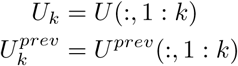

Note:

#### 4.3.4 Self-regularization *f*

*M* is projected onto its top *k* PCs to obtain the rank-reduced SR matrix:

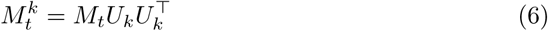

The self-regularization update ((2)) is achieved by taking the difference between the current low- and full-rank SR matrices:

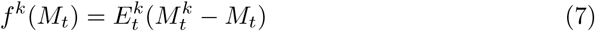

By scaling the difference by the variance explained *E_t_^k^*, the regularization increases dynamically as the low-rank matrix becomes a good representation of the full-rank SR matrix, if it ever does. That is, regularization is proportional to the validity of the low-rank assumption.

*Note on scaling* : One potential feature of scaling by variance explained is to accommodate the agent’s evolving assumptions as it learns. For example, early in learning, evidence for a low-rank structure may not yet be present. Likewise, the full SR matrix *M_t_*would not (yet) have a strong low-rank component, and therefore the influence of the low-rank regularization (7) would be small. This would capture a more complex agent’s evolving strategy in which few assumptions are applied at first (e.g., a full-rank SR is learned with little self-regularization) and then gradually, given the evidence, more assumptions are applied (e.g., learning is guided to a low-rank solution).

Note that *f^k^*(*M_t_*) takes the full matrix *M_t_* and returns a matrix the same size as *M_t_*, but only the row corresponding to the previous state *s_t_*_−1_ is used to update *M_t_*_+1_ (2).

#### 4.3.5 Schema transfer *g*

Schema transfer is applied when the agent has prior experience with the task (i.e., session *>* 1) and therefore has learned the SR matrix *M^prev^*, which is taken as the final state of the SR matrix from the prior session. (For session 1, for which no prior experience exists, no schema transfer is applied, i.e., *ω_prev_*= 0.) Between sessions, the stimuli change, as does the mapping of stimulus categories (i.e., faces, scenes, etc.) to specific states. Therefore, an SR matrix from a prior session cannot be applied verbatim to the current session. However, for any two SR matrices learned on the same meta-structure, we assume their eigenvalues are equivalent (that is, they have the same dimensionality) and that their eigenvectors are equivalent given some rotation (i.e., some permutation of the rows and columns). Furthermore, we assume that the meaningful structure is low-dimensional, and therefore we find the rotation matrix *R* that aligns the top *k* PCs (i.e., first *k* columns) between *M_t_* and *M^prev^*:

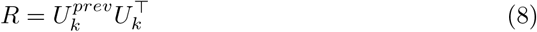

where *U^prev^*are the top *k* PCs of *M^prev^*. (Note that one could instead attempt to align the raw SR matrices directly, but this process would be sensitive to high-dimensional noise, particularly while *M_t_* is early in learning.)

We now define the target matrix for regularization as the component of *M* that overlaps with the first *k* dimensions of *M^prev^*. Similar to above, we apply dimensionality-reduction to the current SR matrix *M_t_* (rotated to the space of *U_k_^prev^*), but now by projecting onto the top *k* PCs from the *previous-session* SR matrix *M^prev^*, and finally rotate back to the space of the current SR matrix:

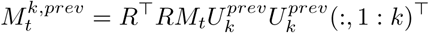

By substituting for *R* (via (8)), this simplifies to:

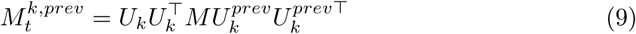

The intuition is that *M_t_^k,prev^* reflects the extent to which *M_t_* represents the low-dimensional components of the “true” SR matrix, which we assume may be better approximated by the well-learned SR matrix from the prior session than the current SR matrix, particularly early in learning. We will then use *M_t_^k,prev^* to regularize *M_t_*.

We also considered applying *U_k_^prev^* symmetrically to the rows and columns:

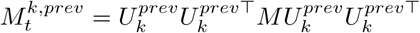

However, this approach consistently fit the data less well, in the log-likelihood sense, than in (9).

The schema transfer update ((3)) is achieved by taking the difference between the target (rotated, low-rank) matrix from the previous session and the current SR matrix:

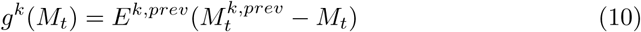

Again, by scaling the difference by the variance explained *E^k,prev^* (in this case, of the low-rank, previous-session matrix), the regularization is weighted by the appropriateness of the low-rank assumption. (Note that, unlike self-regularization, the scaling for schema transfer is *not* dynamic because *M^prev^*, and thereby *E^k,prev^*, are static within a given session.)

Like the self-regularizing function *f* (7), *g* takes the full matrix *M_t_* and returns a matrix of the same size. But *unlike f* , all rows from the output of *g* are used to update *M_t_*_+1_ (3).

#### 4.3.6 Model fitting

The model’s five free parameters {*α, β, γ, ω_self_ , ω_prev_*} were fit to the binary choices from each participant in each session independently using maximum likelihood estimation (i.e., maximizing the likelihood of the data given the model parameters and the estimated probability of choices given by (1)), implemented via MATLAB’s *fmincon* routine. Model parameters were constrained to their respective supports (see above).

Parameter estimation was repeated 20 times independently for each participant-session. For each iteration, parameters were initialized with random values according to their respective supports: *U* (0, 1) for support [0, 1], or *Gamma*(shape = 1, scale = 2) for support [0, ∞]. Parameter estimates from the iteration with the greatest likelihood were selected for further analysis.

Note that while sessions for a given participant were fit independently, the SR matrix at the end of the prior session was used to define *M_prev_* for the current session (see 4.3.5).

Invalid trials were removed prior to model fitting and any subsequent evaluation. The rationale was that, on invalid trials, participants did not render a choice nor learn the outcome of the trial and therefore received no information regarding the state identity. As such, they did not experience a transition from the prior state to the undefined state on the present (invalid) trial and therefore could not update their beliefs about the relationship between the pair of states. Rather, the only defined transition the participant experienced was between the prior state and the state on the next valid trial. By omitting invalid trials, the sequence of trials included for analysis reflected this experience.

#### 4.3.7 Additional notes

##### On selection of rank k

We found that *k* = 1 produced equivalent or better model fits in a log-likelihood sense than higher ranks (*k* ≤ 7) and therefore was used throughout. In the present experiment, with *k* = 1, the low-rank SR resembles a block-diagonal matrix clustering states from the same context. We also tested using unequal ranks for self-regularization *f^k^* and schema transfer *g^l^*^≠*k*^, but no combination of ranks produced better model fits than *k* = *l* = 1 for both functions.

##### On undefined PCs

The PCs of *M_t_* may not be well-defined, particularly early in learning. That is, one or more eigenvalues may be equal, and therefore the set of ordered PCs may non-unique (e.g., on trial 1, *M_t_*_=1_ is set to the identity matrix). In this case, the choice of the top *k* PCs is arbitrary and subject to variation across different systems and/or implementations. To mitigate this variation, self-regularization *f^k^* and schema transfer *g^k^* were not applied on trials 1 and 2 (note that *M_t_* is first updated at the *end* of trial 2, and therefore remains the identity matrix until trial 3). In the present task, this was sufficient to avoid cases of undefined PCs. However, in cases where a higher rank is used (i.e., *k >* 1), additional procedures may be required.

### 4.4 fMRI acquisition

MRI data were collected at the Zuckerman Institute of Columbia University on a 3 T Siemens Magnetom Prisma scanner with a 64-channel head coil. Functional images were acquired using a multiband echo-planer imaging (EPI) sequence (repetition time = 1.55 s, echo time = 30 ms, flip angle = 67°, acceleration factor = 3, parallel reduction factor in plane (iPAT) = 2, phase resolution = 66%, voxel size = 1.71 x 1.71 x 1.70 mm, acquisition matrix 112 × 112). Seventy eight oblique axial slices were acquired in an interleaved order and spaced 1.70 mm to achieve full brain coverage. Whole-brain high resolution (1 mm iso) T1-weighted structural images were acquired with a magnetization-prepared rapid acquisition gradient-echo (MPRAGE) sequence. To estimate field inhomogeneity and aid registration, two *fieldmaps* were collected using the same voxel size, acquisition matrix, and slice number and orientation as the functional scans.

### 4.5 fMRI analysis

#### 4.5.1 Preprocessing

Results included in this manuscript come from preprocessing performed using *fMRIPrep* 24.1.1 [39], which is based on *Nipype* 1.8.6 [40].

##### Preprocessing of B0 inhomogeneity mappings

A *B_0_*-nonuniformity map (i.e., *fieldmap*) was estimated based on two echo-planar imaging (EPI) references with opposite anterior-posterior phase-encoding polarities (i.e., PEPolar) using `topup` [41]. For the one participant without PEPolar acquisitions, the *fieldmap* was estimated from the phase-drift map(s) measure with two consecutive GRE (gradient-recalled echo) acquisitions. The corresponding phase-map(s) were phase-unwrapped using `prelude`.

##### Anatomical data preprocessing

A total of 1 T1-weighted (T1w) images were found within the input BIDS dataset. The T1w image was corrected for intensity non-uniformity (INU) with `N4BiasFieldCorrection` [42], distributed with ANTs 2.5.3 [43], and used as T1w-reference throughout the workflow. The T1w-reference was then skull-stripped with a *Nipype* implementation of the `antsBrainExtraction.sh` workflow (from ANTs), using OASIS30ANTs as target template. Brain tissue segmentation of cerebrospinal fluid (CSF), white-matter (WM) and gray-matter (GM) was performed on the brain-extracted T1w using `fast` [44]. Brain surfaces were reconstructed using `recon-all` [FreeSurfer 7.3.2, 45], and the brain mask estimated previously was refined with a custom variation of the method to reconcile ANTs-derived and FreeSurfer-derived segmentations of the cortical gray-matter of `Mindboggle` [46]. Volume-based spatial normalization to one standard space (MNI152NLin2009cAsym) was performed through nonlinear registration with `antsRegistration` (ANTs 2.5.3), using brain-extracted versions of both T1w reference and the T1w template. The following template was were selected for spatial normalization and accessed with *Template-Flow* [24.2.0, 47]: *ICBM 152 Nonlinear Asymmetrical template version 2009c* [48, TemplateFlow ID: MNI152NLin2009cAsym].

##### Functional data preprocessing

For each of the 8 BOLD runs found per participant (two runs for each of four sessions), the following preprocessing was performed. First, a reference volume was generated, using a custom methodology of *fMRIPrep*, for use in head motion correction. Head-motion parameters with respect to the BOLD reference (transformation matrices, and six corresponding rotation and translation parameters) were estimated before any spa-tiotemporal filtering using `mcflirt` [49]. The estimated *fieldmap* was then aligned with rigid-registration to the target EPI (echo-planar imaging) reference run. The field coefficients were mapped on to the reference EPI using the transform. The BOLD reference was then co-registered to the T1w reference using `bbregister` (FreeSurfer) which implements boundary-based registration [50]. Co-registration was configured with six degrees of freedom. Several confounding time-series were calculated based on the *preprocessed BOLD*: framewise displacement (FD), DVARS and three region-wise global signals. FD was computed using two formulations following Power [absolute sum of relative motions 51] and Jenkinson [relative root mean square displacement between affines 49]. FD and DVARS are calculated for each functional run, both using their implementations in *Nipype* [following the definitions by 51]. The three global signals are extracted within the CSF, the WM, and the whole-brain masks. Additionally, a set of physiological regressors were extracted to allow for component-based noise correction [*CompCor*, 52]. Principal components are estimated after high-pass filtering the *preprocessed BOLD* time-series (using a discrete cosine filter with 128s cut-off) for the anatomical *CompCor* (aCompCor). For aCompCor, three probabilistic masks (CSF, WM and combined CSF+WM) are generated in anatomical space. The implementation differs from that of [52] in that instead of eroding the masks by two pixels on BOLD space, a mask of pixels that likely contain a volume fraction of GM is subtracted from the aCompCor masks. This mask is obtained by dilating a GM mask extracted from the FreeSurfer’s *aseg* segmentation, and it ensures components are not extracted from voxels containing a minimal fraction of GM. Finally, these masks are resampled into BOLD space and binarized by thresholding at 0.99 (as in the original implementation). Components are also calculated separately within the WM and CSF masks. For each CompCor decomposition, the *k* components with the largest singular values are retained, such that the retained components’ time series are sufficient to explain 50 percent of variance across the nuisance mask (CSF, WM, combined, or temporal). The remaining components are dropped from consideration. The head-motion estimates calculated in the correction step were also placed within the corresponding confounds file. All resamplings can be performed with *a single interpolation step* by composing all the pertinent transformations (i.e. head-motion transform matrices, susceptibility distortion correction when available, and co-registrations to anatomical and output spaces). Gridded (volumetric) resamplings were performed using `nitransforms`, configured with cubic B-spline interpolation. Many internal operations of *fMRIPrep* use *Nilearn* 0.10.4 [53], mostly within the functional processing workflow. For more details of the pipeline, see the section corresponding to workflows in *fMRIPrep*’s documentation.

#### 4.5.2 Regions of interest

We defined bilateral anatomical regions of interest (ROIs) *a priori* based on their previously documented contributions to relational learning and abstraction (see 1). Subcortical ROIs—hippocampus and amygdala—were defined using their respective labels in the Harvard-Oxford Subcortical Structural Atlas [54] mapped to the standard space (MNI152NLin2009cAsym) using *TemplateFlow* [24.2.0, 47] with the probability threshold set to 0.

Cortical ROIs were defined using the Julich-Brain Atlas [55] mapped to the standard space using *siibra* [1.0.1-alpha.8, 56] as follows. The entorhinal cortex (EC) ROI was defined as region “Area EC (Hippocampal Region, Entorhinal Cortex)” with the probability threshold set to 0.25. The remaining ROIs were constructed as a union of subregions with `maptype` set to “labelled” (i.e., voxels uniquely assigned to region with the maximum probability). The orbitofrontal cortex (OFC) ROI was created by combining the medial (Areas Fo1, Fo2, and Fo3) and lateral (Areas Fo4, Fo5, Fo6, and Fo7) OFC subregions. The dorsolateral prefrontal cortex (dlPFC) ROI was created by combining subregions of the superior frontal gyrus (SFG2, SFG3, and SFG4) and middle frontal gyrus (MFG4 and MFG5), corresponding to parts of Brodmann areas 9 and 46 [57]. The anterior cingulate cortex (ACC) ROI was defined as region “p24c”, which corresponded to Brodmann area 24c, the site of single-unit recordings in previous, related experiments in monkey [1]. The monkey work focused explicitly on the ventral bank of the cingulate sulcus, whereas our ROI includes the ventral bank and a portion of the fundus and dorsal bank. We believed this extension was reasonable to minimize error in mapping homologous structures between species. Of note, the Julich-Brain region “p24c” refers to the *p*osterior portion of area 24 within the ACC—not to be confused with Brodmann area “p24” of the midcingulate cortex—in contrast to the *s*uperior and rostral Julich-Brain region “s24”, which was subgenual and largely excluded the cingulate sulcus, and therefore not homologous to the recording sites in the monkey [58] .

#### 4.5.3 Statistical model for fMRI analysis

##### Regressors

We conducted a univariate generalized linear model (GLM) to relate BOLD activity to the teaching signals derived from the gSR model. We defined three pairs of regressors of interest, where each pair corresponded to one of the gSR prediction error terms and contained separate regressors for correct and error trials, for a total of six regressors, only three of which were non-zero for a given trial (depending on whether the outcome was correct or error). The magnitude of each regressor corresponded to the trial-specific magnitude of its respective prediction error. Specifically, for the terms corresponding to TD learning and self-regularization, which were in vector form (see (2)), we computed the *L*2 norm, and for the schema transfer term, which was in matrix form (see (3)), we computed the Frobenius norm. Each term was then scaled by its respective free parameter (*α*, *ω_self_* , and *ω_prev_*), as fit for the specific participant and session. Finally, for each term, the trial-wise values were *z* -score normalized by their mean and standard deviation taken over all sessions, facilitating comparisons across sessions. We modeled the error terms as boxcar regressors occurring during the 0.5 s outcome period on each trial.

In addition, we included 12 nuisance regressors. To account for the baseline effect of the outcome, we included two separate binary regressors for correct and error trials, modeled during the outcome period (i.e., same onset and duration as the regressors of interest). To account for variance related to the specific task state (i.e., unique combination of stimulus, optimal action, and outcome value), we included eight binary regressors, one per state, modeled at the onset of the stimulus presentation and with duration set to the median reaction time (RT) for the given participant and session. In addition, we included a single regressor with the same onset and duration but modulated by the trial-specific RT. Lastly, we included a binary regressor on invalid trials, modeled from the onset of the stimulus presentation to the time the outcome period would have terminated had the participant provided a valid response.

Finally, we included 18 confound regressors to control for movement, physiological, and signal artifacts, as generated by *fMRIPrep*: framewise displacement, six *CompCor* regressors (i.e., first six principal components of activity in a combined CSF and white matter mask), six rigid-body motion parameters (three translations and three rotations), and a set of five discrete cosine transform basis functions to control for low-frequency signal drifts [24.1.1, 39].

##### GLM model estimation and hypothesis testing

Model estimation and hypothesis testing were performed with FSL’s FEAT [6.0.4, 54]. The first-level time-series GLM analysis was estimated for each run per participant (parameters: spatial smoothing 5mm FWHM, FILM prewhitening, no high-pass temporal filtering). The resulting coefficients (*a*, *b*, *c*) for the regressors of interest (i.e., gSR prediction error terms) were entered into one or more linear contrasts that compared the coefficient(s): *a* to baseline (i.e., whether BOLD activity was modulated by the corresponding regressor above and beyond that accounted for by the remaining regressors in the GLM), *a* − *b* (i.e., whether BOLD activity was modulated by one regressor of interest more than another), and *a* − (*b* + *c*) (i.e., whether BOLD activity was modulated by one regressor of interest more than the combination of both other regressors of interest). Each of these contrasts was computed both for the regressors corresponding to correct trials only (i.e., *a* = *a_correct_*) and by combining correct and error trials (i.e., *a* = *a_correct_* + *a_error_*). Given three gSR terms, this produced a total of 18 (i.e., (3 + 3 + 3) ∗ 2) contrasts.

The first-level contrasts were then combined across runs per participant using fixed effects. Because the gSR schema transfer term was not defined for session 1 (no prior session existed from which to generalize), we restricted analysis to runs from sessions 2, 3 and 4, which were combined in each of two ways. First, to test for an overall effect across sessions (“overall”), each run was weighted equally (i.e., scaled by 1 for each of sessions 2, 3, and 4). Second, to test for a linear trend across sessions (“increasing” or “decreasing”), runs were weighted proportional to their session number (i.e., scaled by -0.5, 0, or 0.5 for sessions 2, 3, or 4, respectively).

The second-level, across-session contrasts were combined across participants using FSL’s mixed effects modeling tool FLAME1 and normalized to z-statistics [59] Group-level maps were corrected to control the familywise error rate via cluster-based Gaussian random field correction for multiple comparisons, with an uncorrected cluster-forming threshold of z-statistic = 2.3 and corrected extent threshold of p ¡ 0.05. This correction procedure was performed separately within each predefined ROI (see 4.5.2).

### 4.6 Post-hoc exclusions

We excluded participants post-hoc for inattention during the behavioral task based on two features. First, we excluded participants with unusually poor performance on non-inference trials (i.e., trials on which optimal performance depended only on trial-and-error learning and did not require relational knowledge of the task structure).

Specifically, we examined performance on the third and subsequent repetitions of each stimulus within each block and flagged sessions in which accuracy on these trials was at or below 80%. We excluded participants with two or more flagged sessions. Five participants were excluded for performance-based inattention, all of whom had below-threshold accuracy on all four sessions, typically in the range of 45 - 70%.

Second, we excluded participants with a high number invalid responses (e.g., not responding in time), which impacted the reliability of both the gSR model fits and the subsequent fMRI GLM estimation. Specifically, we excluded participants with an average of more than five invalid trials per session or more than 10 invalid trials in two or more sessions. A single participant met both criteria and was excluded.

## Notes

### Competing Interest Statement

The authors have declared no competing interest.

### Summary of Updates

The author affiliations were updated and minor typographical errors were corrected.

